# Computational 4D-OCM for label-free imaging of collective cell invasion and force-mediated deformations in collagen

**DOI:** 10.1101/2020.09.10.291633

**Authors:** Jeffrey A. Mulligan, Lu Ling, Claudia Fischbach, Steven G. Adie

## Abstract

Traction force microscopy (TFM) is an important family of techniques used to measure and study the role of cellular traction forces (CTFs) associated with many biological processes. However, current standard TFM methods rely on imaging techniques that do not provide the experimental capabilities necessary to study CTFs within 3D collective and dynamic systems embedded within optically scattering media. Traction force optical coherence microscopy (TF-OCM) was developed to address these needs, but has only been demonstrated for the study of isolated cells embedded within optically clear media. Here, we present computational 4D-OCM methods that enable the study of dynamic invasion behavior of large tumor spheroids embedded in collagen. Our multi-day, time-lapse imaging data provided detailed visualizations of evolving spheroid morphology, collagen degradation, and collagen deformation, all using label-free scattering contrast. These capabilities, which provided insights into how stromal cells affect cancer progression, significantly expand access to critical data about biophysical interactions of cells with their environment, and lay the foundation for future efforts toward volumetric, time-lapse reconstructions of collective CTFs with TF-OCM.

## Introduction

Cellular traction forces (CTFs) play a critical role in both physiological and pathological tissue remodeling processes, including angiogenesis^1,2^, collective migration^3–5^, and cancer metastasis^6,7^, and can vary as a function of extracellular matrix (ECM) remodeling and cellular phenotype. For example, compositional or structural changes of the ECM change the CTFs that cells exert upon the surrounding matrix^8^, and these changes in CTFs, in turn, alter cellular mechanosignaling, which reciprocally modulates CTFs^9^. Traction force microscopy (TFM) is a diverse family of techniques used to quantify and study the CTFs that cells exert upon the ECM^10–12^. Specifically, TFM reconstructs CTFs from optical measurements of CTF-induced ECM deformations using ECM rheological data and a suitable mechanical model. By providing measurements of both CTFs and CTF-induced ECM deformations, TFM serves as a vital tool for studying the biophysical interactions of cells with the ECM (e.g., forces, degradation, and remodeling) and the role that these interactions play within the broader field of mechanobiology^13^. Consequently, there is an ongoing demand for the development of novel TFM capabilities that satisfy the evolving needs of mechanobiology to investigate CTF dynamics within complex ECM substrates.

There are substantial gaps between the capabilities of current standard TFM technologies and modern research needs. For example, the vast majority of TFM techniques are limited to the study of 2D cell culture models^14–19^. However, since cells can exhibit drastically different phenotypes in 2D versus 3D environments^13,20–23^, 3D TFM techniques are crucial for studying cells under physiologically relevant 3D conditions. Although 3D TFM methods have been developed in response to this need^24–28^, the confocal microscopy paradigm which these techniques rely upon restricts their capabilities. In particular, a limited penetration depth (of a few hundred micrometers in optically scattering media) hinders the study of either deeply embedded cells or extended multicellular collectives within scattering substrates. Photobleaching and/or phototoxicity impede long-term observations of dynamic interactions and feedback between CTFs and ECM remodeling (and thereby limit 3D TFM studies to endpoint observations and/or brief/sparse time-sequences)^18,28^. These concurrent limitations have culminated in a lack of readily available TFM technologies which simultaneously enable the study of 3D, collective, and dynamic cell behaviors within optically scattering media. As a consequence, TFM research continues to employ cell culture models (e.g., 2D instead of 3D) which can differ markedly from the systems that researchers seek to understand^5^. This has created a significant opportunity for new TFM techniques to enable experiments that are impractical or impossible with current standard methods.

In order to meet the growing demand for TFM methods which enable the study of 3D, collective, and dynamic behaviors, we recently developed traction force optical coherence microscopy (TF-OCM)^29,30^. This technique leverages optical coherence microscopy (OCM) to provide rapid, label-free, volumetric imaging within optically scattering media. As a result, the technique is well-suited to both rapid and long-term 3D time-lapse imaging with numerous exposures (since photobleaching and photoxicity are not of concern due to a lack of fluorescent labels and a low incident power, <5 mW). Furthermore, the use of near-infrared light and coherence gating enables imaging deep within scattering media, such as collagen or other biopolymer substrates (up to 1 millimeter and deeper). As a result, TF-OCM is a promising candidate to perform studies of 3D, collective, and dynamic systems within physiologically relevant substrates. However, TF-OCM has only been demonstrated for the study of isolated cells within optically clear (Matrigel) substrates^29,30^.

Here, we report on the use of 4D-OCM imaging and data processing methods (henceforth referred to as ‘computational 4D-OCM’) to study the invasion of tumor spheroids embedded within optically scattering 3D collagen substrates over long periods of time. Spheroids consisting of premalignant tumor cells and/or stromal cells were fabricated and embedded into dense collagen matrices to mimic cellular interactions with a surrounding fibrotic ECM characteristic of tumors. Subsequently, collective tumor cell and/or stromal cell invasion into the surrounding collagen and corresponding ECM remodeling were monitored over 2 days using a 40-minute imaging interval. In order to obtain mm^3^-scale volumetric images with high spatial resolution and minimal distortions, we developed and employed a novel OCM system design in combination with a refined set of numerical image formation techniques which together provided the spatial coverage and resolution required for these new experimental conditions. Importantly, by leveraging the label-free and long-term imaging capabilities of our OCM system, we were able to measure time-varying 3D displacement fields within the collagen substrate(s) *without* the use of either fiducial marker beads or contractility inhibiting reagents, since such additives (although ubiquitous in standard TFM methods) may alter cell behavior and/or preclude long-term, continuous observation of individual spheroid systems. One example of the potential impact of the methods reported here can be found in our recently published study^31^ that investigated the role of stromal cells in CTF-dependent collective tumor cell invasion under clinically relevant conditions. As a result, computational 4D-OCM has been demonstrated to be a promising technique for the study of biophysical cell-ECM interactions and collective cell behavior in physiologically relevant model systems. These developments provide a critical foundation for enabling future work toward time-lapse reconstructions of collective CTFs with TF-OCM.

## Results

### Overview of experiments, context, and findings

In this study, spheroids (consisting of 5-10×10^3^ cells) were embedded in dense 3D collagen matrices. Upon completion of this embedding, time-lapse imaging was begun immediately in order to monitor all subsequent spheroid invasion and collagen deformation/remodeling via our computational 4D-OCM imaging methods (detailed in the Methods). Imaging was performed at 40-minute intervals for a total duration of 2 days, after which custom data processing routines (detailed in the Methods and Supplementary Note 3) were used to reconstruct 4D image data and visualize/quantify the biophysical interactions between cell collectives and 3D collagen substrates. Our computational 4D-OCM methods (and subsequent data visualizations) for imaging cell invasion and collagen deformation/remodeling are detailed in the remaining sections of this manuscript, whereas biological findings arising from the application of these methods to study the role of obesity-associated adipose stromal cells (ASCs) in cancer progression can be found in a companion paper^31^.

Computational 4D-OCM imaging of biophysical cell-ECM interactions was used to study multiple spheroids of differing compositions, consisting of monocultures and/or co-cultures of both tumor cells and/or ASCs. In particular, tumor cells were modeled using MCF10AT1 cells, a premalignant human breast epithelial cell line. Tumor spheroids consisting of these cells (in monoculture) do not typically invade dense collagen matrices such as those used here^31^. ASCs, on the other hand, can be used to form highly invasive spheroids (e.g., as in Fig. 1) and have previously been shown to promote malignant behavior in tumor cells, including increased proliferation and migration^32^. In this study, two types of (murine) ASCs were used, derived from subcutaneous fat of both lean (wildtype, WT) and obese (*ob/ob*) mice, respectively. In Ref. ^31^, it was shown (using our computational 4D-OCM methods) that while both WT and *ob/ob* ASCs were highly motile/invasive, *ob/ob* ASCs induced greater ECM deformations both in monoculture and co-culture (with MCF10AT1 cells) than their WT counterparts. (Please refer to Ref. ^31^ for additional data and quantitative comparisons regarding these findings.)

**Figure 1.**
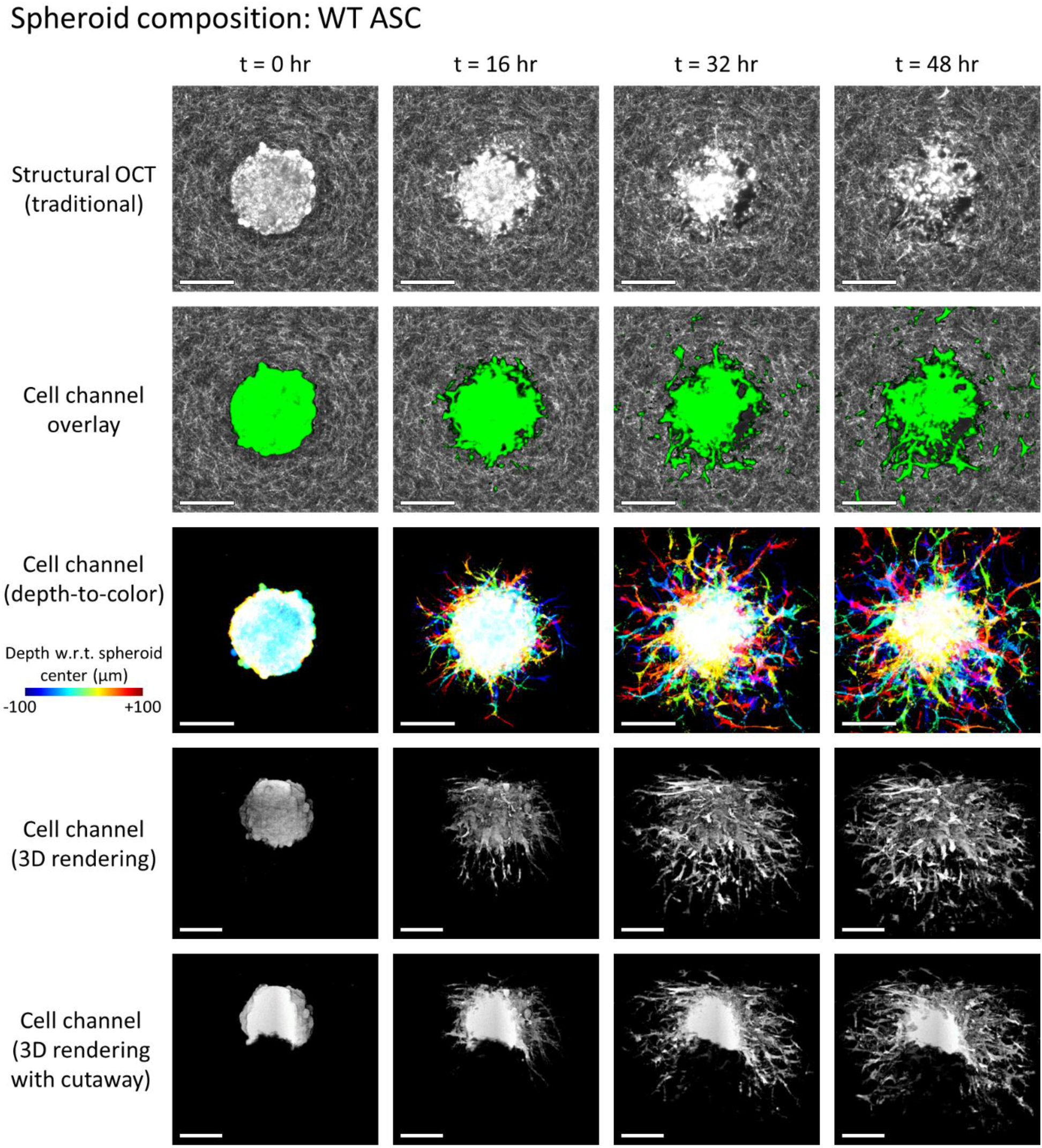
Invasion of a WT ASC monoculture spheroid into the surrounding collagen substrate, revealed via label-free OCM imaging and temporal speckle contrast. Traditional OCM imaging (top row) records scattering signals of cells and collagen alike. Temporal speckle contrast enables segmentation of volumetric data into synthetic ‘cell’ and ‘collagen’ channels (rows 2-5). Scale bars = 200 μm. See text for details. Time-lapse animation is available in Supplementary Video 1.

In the sections that follow, we highlight the imaging capabilities provided by our computational 4D-OCM methods and provide details regarding the underlying imaging methods and data processing routines. In particular, we demonstrate the 4D label-free visualization of spheroids and their invasion into the surrounding collagen (Figs. 1 and 2). This visualization is greatly enhanced by the segmentation of cells from the collagen substrate, which was achieved using only temporal fluctuations of backscattered optical signals. We further visualize remodeling of the collagen substrate, in the form of both degradation (Fig. 3) and CTF-induced deformation (Fig. 4), without the addition of any labels or fiducial marker beads to the substrate medium. Together, these data, which could not otherwise be obtained through more conventional confocal fluorescence and/or reflectance microscopy-based techniques, were used to make essential contributions to the study detailed in Ref. ^31^. Specifically, quantitative comparisons of differing spheroid invasion and ECM remodeling behaviors (not shown here) may be found in the manuscript for that study^31^. These findings and contributions demonstrate the utility of computational 4D-OCM for making novel contributions to mechanobiology research.

**Figure 2.**
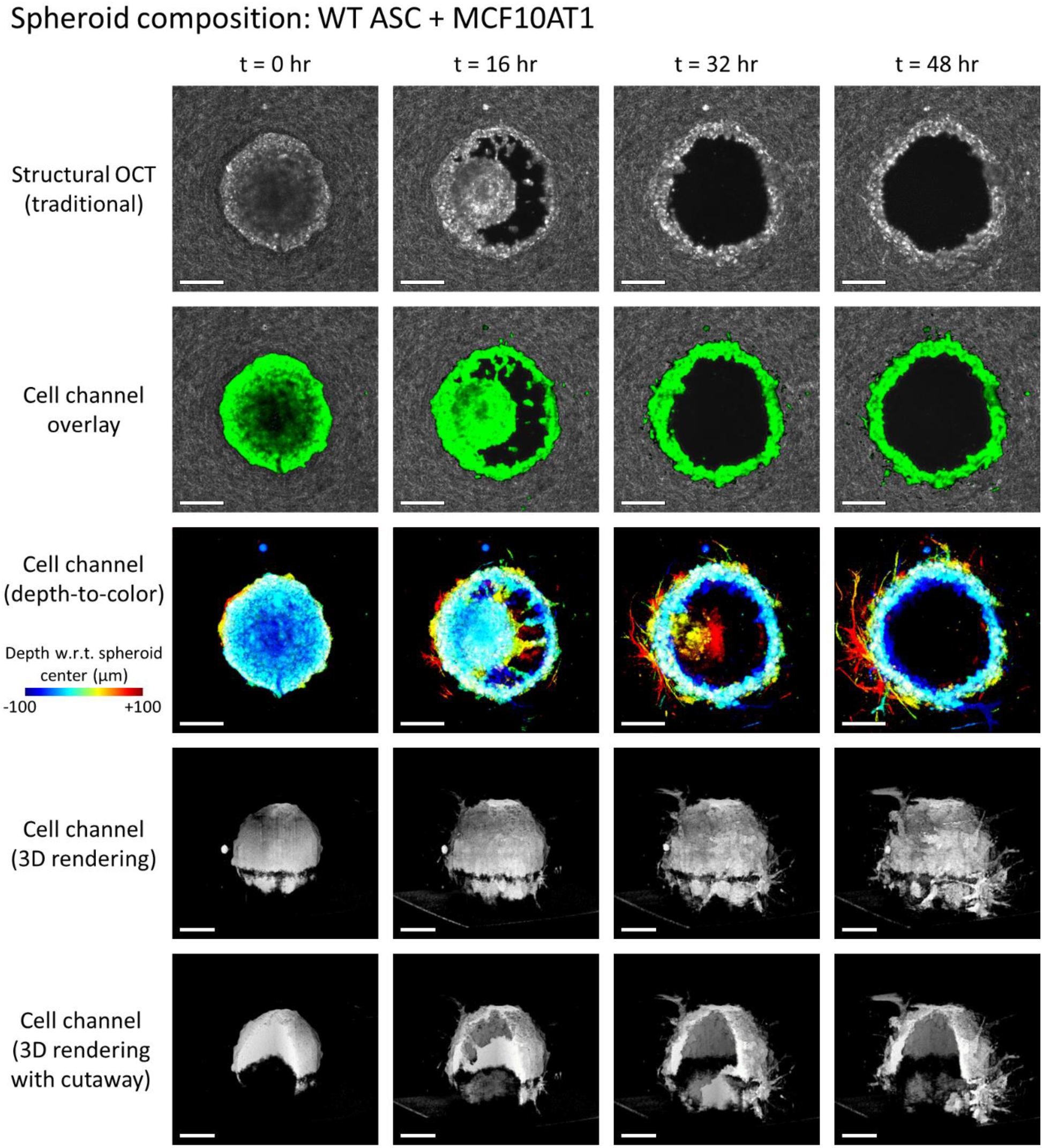
Invasion of a WT ASC + MCF10AT1 co-culture spheroid into the surrounding collagen substrate, revealed via label-free OCM imaging and temporal speckle contrast. Traditional OCM imaging (top row) records scattering signals of cells and collagen alike. Temporal speckle contrast enables segmentation of volumetric data into synthetic ‘cell’ and ‘collagen’ channels (rows 2-5). Scale bars = 200 μm. See text for details. Time-lapse animation is available in Supplementary Video 2.

**Figure 3.**
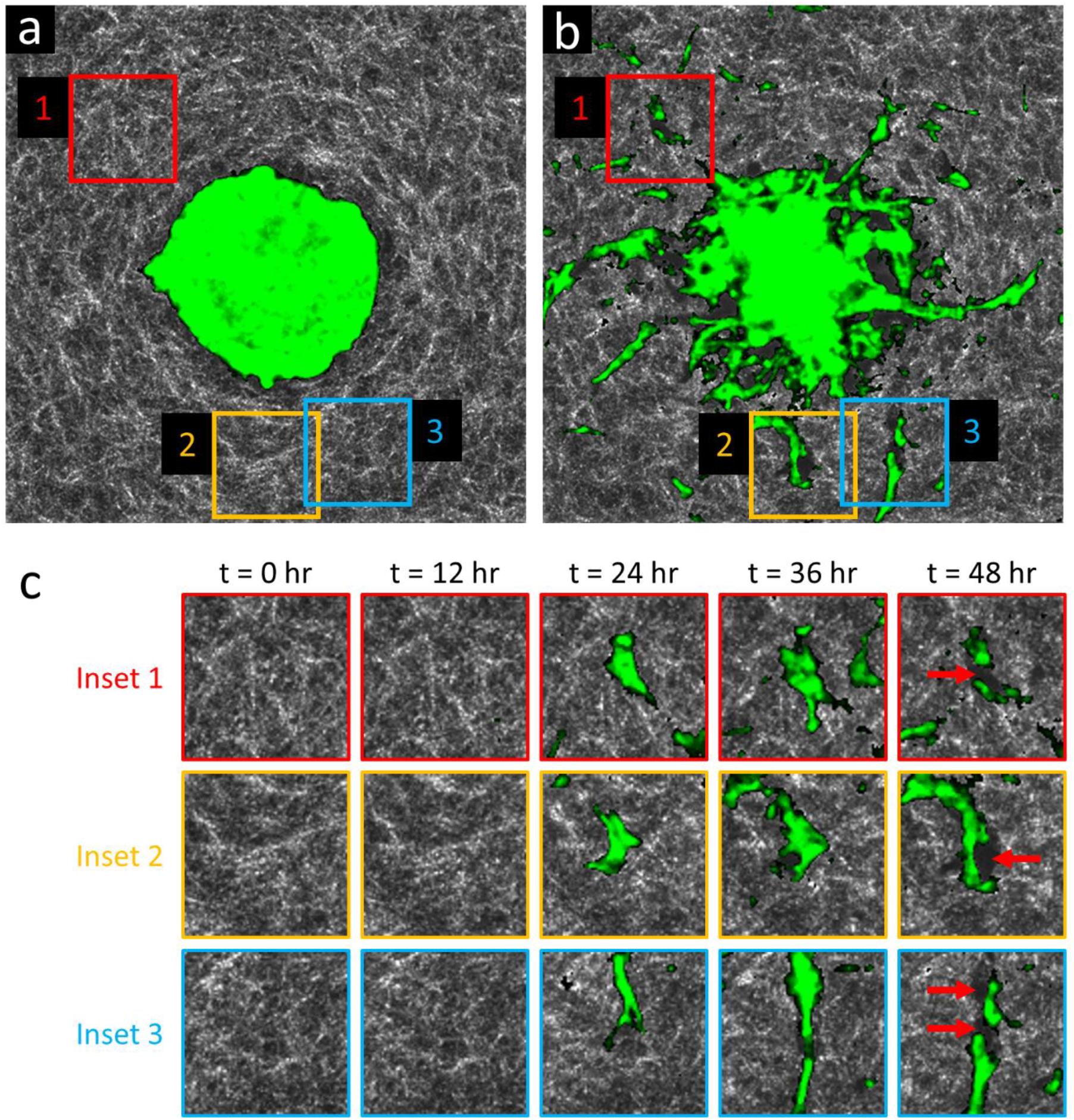
Time-lapse OCM imaging reveals collagen degradation by invasive cells and strands of cells. (a) *En face* plane intersecting the center (i.e., the ‘equator’) of a lean (WT) ASC monoculture spheroid, acquired at time t = 0 hr. The spheroid (green, highlighted via temporal speckle contrast) is recently embedded, and has not yet invaded the surrounding collagen (white). (b) The same *en face* plane as in (a), acquired at time t = 48 hr. Invasive protrusions are abundant. Large dark regions surrounding the spheroid and invasive strands correspond to ‘voids’ with low/weak scattering signals, suggesting a lack of either cells or collagen. (c) Time-lapse view of insets 1-3 from (a,b). Red arrows at time t = 48 hr indicate newly formed ‘void’ regions where only collagen was initially present. These new ‘voids’ are likely due to degradation of the collagen matrix by invasive strands via matrix metalloproteinase activity. Panels (a,b) span a 750×750 μm^2^ lateral FOV. Time-lapse animation is available in Supplementary Video 3.

**Figure 4.**
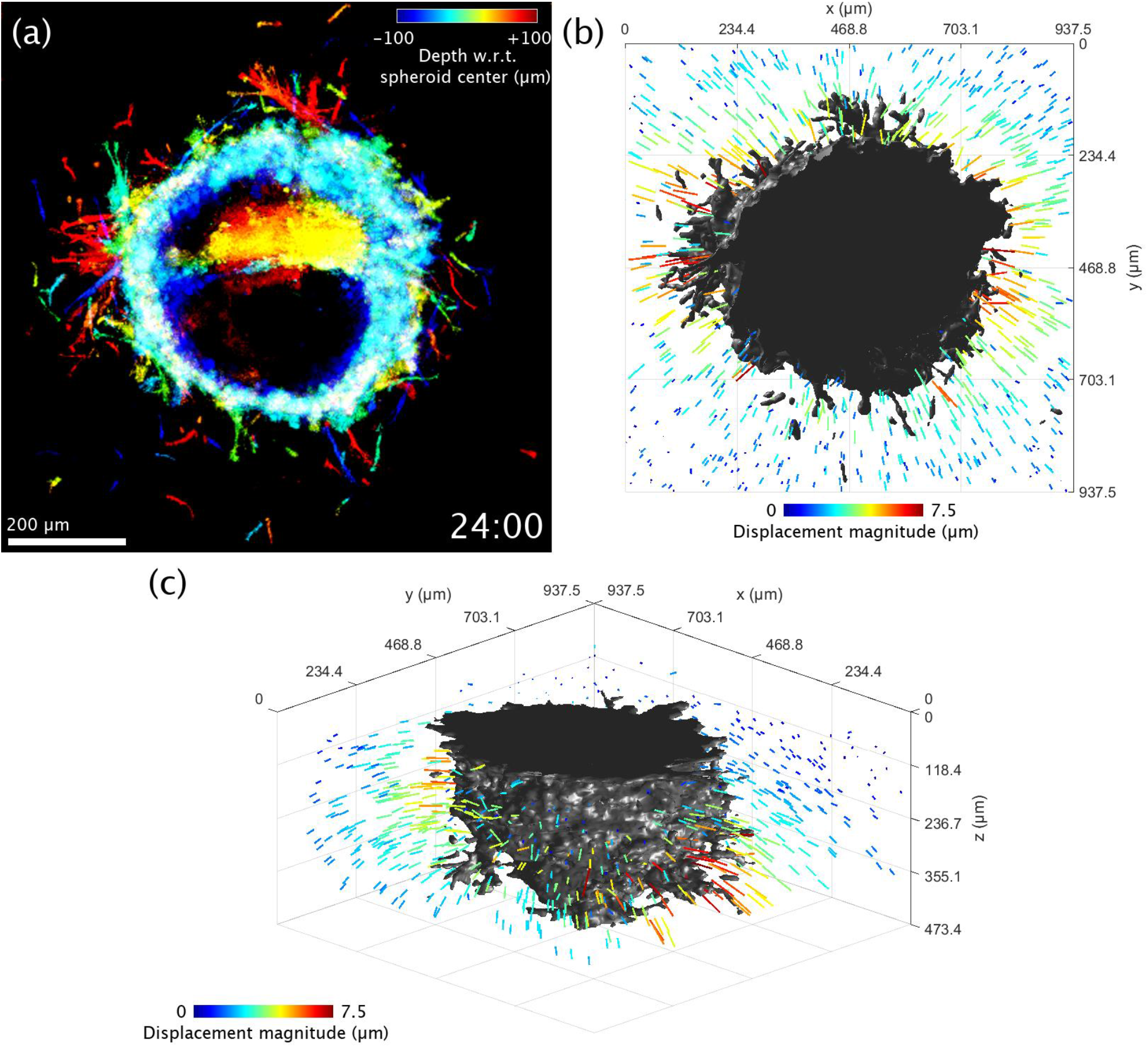
Visualization of 3D collagen displacements in the vicinity of an obese (*ob/ob*) ASC + MCF10AT1 co-culture spheroid at time t=24 hr. (a) Top-down view of the spheroid, using the same depth-to-color projection as in row 3 of Figs. 1 and 2. (b) Top-down rendering of the spheroid (shown in gray) accompanied by colored arrows which indicate displacement of the collagen matrix as measured with respect to its initial configuration at time t=0 hr. Arrow lengths have been exaggerated for visualization purposes. (c) Re-rendering of panel (b) from an isometric viewing angle. The flat surface of the spheroid corresponds to a region where the spheroid comes into contact with the coverslip bottom of the petri dish. Time-lapse animation is available in Supplementary Video 4.

### Temporal speckle contrast enables label-free, 4D visualization of spheroid invasion behavior in collagen

One challenge of using label-free imaging (as is the case with OCM) is to identify the boundary between cells and the surrounding collagen substrate. In order to address this challenge, we leveraged a ‘burst’ imaging protocol similar to our previously reported TF-OCM methods^30^. For each time-point in a given time-lapse experiment, a ‘burst’ of nine volumetric images was acquired in rapid succession. Due to the short time period (~12-15 seconds) between individual acquisitions in a given ‘burst’, the observed speckle pattern from (quasi-static) collagen fibers remained practically constant. In contrast, the observed speckle pattern from cells exhibited substantial variation between acquisitions, due to the rapid motion of intracellular components. In order to take advantage of this temporal speckle contrast, the images of a given ‘burst’ were combined into a single volumetric image via a ‘standard deviation projection’ operation, as defined in Eqn. (2). The resulting image(s) exhibited strong/enhanced contrast in regions with temporally unstable speckle patterns, and weak/suppressed contrast in regions with temporally stable speckle patterns, consistent with similar existing methods^30,33,34^. This enabled the generation of two synthetic ‘channels’ from otherwise label-free OCM image data: 1) A ‘cell’ channel which strongly correlates to live/active cells and 2) a ‘collagen’ channel which strongly correlates to nonliving/static collagen. (Quotes are used here to signify that the correspondence of a given signal to actual cells and/or collagen lacks the direct specificity of fluorescent labels.)

Representative results of this process are depicted in Fig. 1 and Fig. 2. Row 1 (structural OCT) shows standard label-free OCM images, corresponding to *en face* planes which intersect the center (i.e., the ‘equator’) of the given spheroid. Although it may be easy to distinguish the spheroid from the surrounding collagen by eye at time *t* = 0, the distinction becomes less clear as time (and spheroid invasion) progresses, especially in Fig. 1. However, in row 2 (cell channel overlay), a clear picture of invasion activity is obtained via false-color ‘labeling’ of regions with high temporal speckle contrast. By extension, the 3D structure of the invading spheroids is revealed by the synthetic ‘cell’ channel (rows 3-5). Row 3 depicts a top-down view of this ‘cell’ channel, wherein each depth in the image is assigned a unique color based on its distance from the plane shown in rows 1 and 2, and the final image is obtained via a maximum intensity projection of each color channel along the depth axis. Rows 4 and 5 depict 3D renderings of the ‘cell channel’, where row 4 shows the surface of the spheroid from an isometric viewing angle, and row 5 shows a ‘cutaway’ which reveals the interior structure of the spheroid. The wildtype (WT, lean) ASC monoculture spheroid in Fig. 1 retains a solid/confluent internal structure over time, whereas the WT ASC + MCF10AT1 co-culture spheroid in Fig. 2 exhibits a central ‘core’ of cells which detached from the exterior ‘shell’ and migrated along the depth axis until it was no longer visible to the imaging system. The presence of these distinct structures in the co-culture spheroid is consistent with histology, such as that shown in Fig. S1 of Ref. ^31^.

### Label-free scattering contrast provides visualization of collagen degradation

In addition to the ‘cell’ and ‘collagen’ channels discussed above, we also observed a third feature in the image data: ‘voids’. These regions of low scattering signal correspond to one of two features: 1) areas which cells initially occupied but vacated over time, leaving behind fluid-filled spaces and cavities, or 2) areas in the collagen substrate which have been degraded by invading cells (and are not currently occupied by cells). Both of these features are depicted in Fig. 3, with Fig. 3(c) highlighting regions of newly degraded collagen, in particular. The ability to identify these modifications to the collagen substrate is potentially useful for studying the mechanisms underlying the collective invasion process^31^. Unfortunately, these regions are also (currently) prohibitive to conducting quantitative CTF reconstructions with TF-OCM due to the complicated and heterogeneous mechanical environment + boundary conditions that they present (further details are provided in the Discussion).

### Elastic image registration reveals space- and time-varying collagen displacements

Time-varying deformations of the collagen substrate were observed in conjunction with the previously described changes in spheroid morphology and collagen degradation/remodeling. Unlike standard TFM methods which track the motion of embedded fiducial marker beads, here, the motion of collagen fibers was tracked *directly* (analogous to a previously reported TFM method based on confocal reflectance microscopy^27^). Elastic image registration (via the built-in MATLAB function *imregdemons*) was employed to estimate the time-varying 3D displacement field which would map the original collagen substrate at time *t*=0 to the deformed collagen substrate observed at any given time-point during the first 24 hours of imaging. As a result, the time-varying collagen displacement field could be quantified and visualized, as in Fig. 4 and Supplementary Video 4. (Note that Fig. 4 and Supplementary Video 4 depict collagen displacements measured around an *ob/ob* ASC + MCF10AT1 co-culture spheroid. These spheroids demonstrated the most dramatic displacement fields among all imaged spheroids^31^.) These data revealed contraction of the collagen toward the spheroid body, with the strength of contraction varying as a function of position. Regions of collagen with stronger displacements likely indicate areas where the spheroid may be exerting stronger contractile forces, although a quantitative reconstruction of CTFs would be required to verify this hypothesis^35^. However, even in the absence of CTF reconstructions, this deformation data alone can provide valuable information for learning about cell/spheroid behavior^31,36^.

## Discussion

The results of the previous section demonstrate several advantages provided by our computational 4D-OCM imaging methods. Micrometer-scale details of both spheroid and collagen structure were observed over a large mm^3^-scale volumetric field-of-view using only native scattering contrast. Rapid volumetric image acquisition (on the order of 10-15 seconds per volume, as detailed in the Methods) enabled the measurement of temporal fluctuations in the backscattered optical signals from cells, and these fluctuations were successfully exploited to enable the segmentation of cells from the surrounding collagen. As a result, 3D spheroid morphology was readily extracted from the optically scattering collagen background without the aid of labeling agents. Furthermore, due to the low phototoxicity inherent to OCM imaging, spheroid morphology was readily monitored in detail throughout multi-day time-lapse experiments. Finally, since the scattering contrast provided by OCM also captured time-varying collagen structure, collagen degradation could be observed in the vicinity of invading cellular protrusions, and spatiotemporally heterogeneous deformations of the collagen substrate could be quantified. Unlike standard TFM methods, 4D-OCM was able to track these collagen deformations directly, without the addition of artificial fiducial marker beads. These combined imaging capabilities have enabled access to previously inaccessible data that is critical to advancing mechanobiology research in clinically relevant settings^31^. Taken together, this establishes computational 4D-OCM as a promising experimental technique for 4D imaging of biophysical cell-ECM interactions and collective cell behavior in physiologically relevant 3D collagen substrates.

Temporal speckle contrast has proven remarkably useful for distinguishing invading cells from the surrounding (quasi-static) collagen substrate. However, this method does have weaknesses. In particular, in our previous study^30^, we showed that fine cellular protrusions (e.g., filipodia) can exhibit relatively static speckle patterns over the time-scales of our ‘burst’ imaging protocol (i.e., a volumetric acquisition rate on the order of 0.1 Hz repeated over a total duration of 1.5-2 minutes). We expect similar limitations to be present in this study. As a result, fine/static structures may exhibit weak contrast after performing our ‘standard deviation projection’ operation, and thus be incorrectly placed within the ‘collagen’ channel of our data. Future work with co-registered OCM and confocal fluorescence microscopy images will be required to quantify the severity of this effect. Our use of temporal speckle contrast also substantially increases the amount of raw data that must be acquired by nearly an order of magnitude. However, the ability to generate synthetic imaging channels from label-free, long-term OCM image data may outweigh this cost. Nevertheless, future work which seeks to reduce the amount of data that must be acquired would be of great value. If large quantities of data *are* acceptable, and high-speed OCM imaging is available, the utility of temporal speckle contrast may be further enhanced by enabling the generation of multiple synthetic channels which correspond to different structures and/or levels of metabolic activity^34^.

Although we were able to quantify the displacement of collagen fibers via elastic image registration (using the MATLAB function *imregdemons*), these measurements cannot be expected to be reliable very close to the cell-collagen boundary. This is because *imregdemons* uses Thirion’s ‘Demons algorithm’^37,38^, which performs image registration via an iterative optimization procedure inspired by the optical flow equations. This ill-posed problem is typically regularized via a diffusion (blurring/smoothing) process over the (iteratively updated) displacement field and/or by imposing other constraints (such as requiring that the computed displacement field describes a diffeomorphic transformation)^37^. Unfortunately, since invading cells can degrade and migrate through the 3D collagen substrate, a displacement field which ‘perfectly’ registers collagen fibers over time may exhibit jump discontinuities and/or be non-invertible at the cell-collagen interface. The typical regularization schemes of the Demons algorithm (i.e., isotropic and homogeneous spatial averaging) cannot accommodate these types of features. Algorithms based on local cross-correlation/matching would be similarly vulnerable, due to the spatial averaging inherent to cross-correlation windows. In the event that migrating cells deposit new collagen substrate, a valid displacement field may not even exist at all locations within the collagen substrate at a given time-point. In order to obtain reliable collagen displacement data near the cell-collagen boundary (when such a measurement is possible), alternative algorithms must be found or devised which account for the complex boundary conditions which occur over long-term multicellular invasion processes. For example, regularization of the Demons algorithm via a spatial averaging scheme which ignores cell-occupied regions (via spatially-varying averaging kernels) may be a viable option. If such a solution cannot be found, the incorporation and tracking of scattering fiducial marker beads within the collagen substrate may be a suitable alternative so long as the presence of beads does not appreciably alter cellular behavior. One advantage of the label-free contrast leveraged by our computational 4D-OCM methods is that, unlike fluorescence-based TFM techniques for which marker beads are typically *required*, our results here demonstrate that marker beads are merely *optional* when using computational 4D-OCM. By extension, future variations of TF-OCM built upon this technique may be particularly attractive for studying cells within physiologically relevant 3D collagen substrates.

Long-term imaging with computational 4D-OCM offers another interesting advantage for TF-OCM. In order to understand this advantage, we first require some background knowledge about TFM. In order to measure substrate deformations, most TFM techniques require at least two images (any technique which does *not* require two images must have some form of *a priori* knowledge about the substrate structure^11,39^). These images consist of a ‘reference’ state (i.e., when the substrate is relaxed and no appreciable CTFs are present) and a ‘deformed’ state (i.e., when CTFs are present and the substrate is deformed). In typical TFM applications these two images are acquired as follows: 1) cells are embedded and allowed to achieve a contractile state (this may require several hours in an incubator), 2) an image of the resulting ‘deformed’ state is acquired, 3) a contractility inhibitor or lethal reagent is introduced to induce cell relaxation, and 4) an image of the resulting ‘reference’ state is acquired. This is a relatively rapid and simple procedure. However, it relies on a few important assumptions: 1) the introduced reagent completely eliminates CTFs, 2) the substrate becomes fully relaxed (i.e., with no residual stress due to cell-mediated remodeling of the original substrate or other mechanisms), and 3) no further information about the temporal evolution of the biological system under physiological conditions is required (since physiological conditions and normal cell behavior are disrupted after the addition of force-inhibiting and/or lethal reagents). In this study, since we wished to observe the long-term behavior of invading cells which were free to remodel the collagen substrate, an ‘artificial’ chemically-induced reference state was not acceptable. Instead, by imaging the collagen substrate and spheroid immediately after embedding, we obtained a ‘natural’ reference state before sufficient time had passed for cells to appreciably deform the collagen substrate. (This method is corroborated by a similar ‘natural’ reference state approach that was recently applied to TFM for embedded spheroid systems using far-field deformation measurements^35^.) As a result, we were able to continuously monitor the evolving substrate deformations as long as our image registration algorithm could be trusted. (Here, displacement tracking was halted after 24 hours, since invasions into the collagen became substantial beyond that time.) Long-term imaging experiments with computational 4D-OCM may offer new opportunities to study plastic deformations/collagen remodeling by following a new procedure: 1) acquire the ‘natural’ reference state immediately after spheroid embedding, 2) observe CTF-induced deformations over time, 3) when sufficient time has elapsed, introduce a contractility inhibitor, and 4) acquire the ‘artificial’ reference state. Any discrepancies between the ‘natural’ and ‘artificial’ reference states may offer a means to quantify residual/plastic deformations and/or residual CTFs. We are not aware of any current TFM methods which provide/use such measurements.

The reconstruction of CTFs exerted by large multicellular constructs embedded in collagen over long time-scales is a challenging problem that has yet to be fully addressed by the TFM field^5^. In addition to the challenges of image segmentation and displacement tracking (discussed above), force reconstruction within collagen is a challenging problem in its own right^11,26,40,41^. In particular, collagen is mechanically nonlinear and often exhibits heterogeneity and anisotropy across a range of spatial scales. Moreover, cells can remodel the collagen matrix, altering its mechanical properties over time. This makes collagen supremely difficult to characterize and model to the degree necessary for numerical CTF-reconstruction schemes. Although TFM researchers have begun to make forays into addressing this challenge^26,35^, much work remains in order to enable high-resolution CTF reconstructions for large and dynamic multicellular constructs. In the meantime, our computational 4D-OCM imaging methods provide data that is useful to mechanobiology research (e.g., spheroid/cell morphology and collagen displacement data), even in the absence of quantitative CTF reconstructions^31,36,40^. By making such data more readily available for studies of multicellular systems embedded in scattering media, our methods may further help to provide necessary data to aid ongoing and future research toward achieving quantitative CTF reconstructions. Other OCM-based techniques, such as optical coherence elastography, may also be used alongside our computational 4D-OCM imaging methods to aid in the characterization of local and dynamic mechanical properties within biopolymer substrates^42–45^.

## Conclusion

In this study, our previously developed (single-cell) TF-OCM imaging methods^30^ were adapted to enable the study of the dynamic invasion behavior of tumor spheroids embedded within 3D scattering collagen substrates. By leveraging temporal speckle contrast to distinguish live cells from the background collagen substrate, our methods revealed detailed spheroid morphology throughout multi-day, time-lapse experiments without the use of either endogenous or exogenous labeling. The label-free contrast mechanism underlying OCM imaging further revealed cell-induced collagen remodeling and CTF-induced deformations of the collagen substrate. This was achieved without the use of fiducial marker beads or disruptive contractility inhibiting reagents that are otherwise ubiquitous to TFM. Although much work remains to enable quantitative reconstructions of CTFs within highly nonlinear (and possibly remodeled) collagen substrates, the new capabilities afforded by our current methods nevertheless provided access to valuable data that have already facilitated the advancement of broader mechanobiology studies^31^. Altogether, computational 4D-OCM provides a promising approach to bridge the gap between current experimental TFM capabilities and the needs of mechanobiology researchers to study 3D, collective, and dynamic cell behaviors within optically scattering media. As a result, future research and development toward the quantitative reconstruction of collective cell forces with TF-OCM may enable new and promising investigations of biophysical phenomena.

## Methods

### Animal use and regulatory compliance

All experiments using cells isolated from animals were approved by the Cornell University Institutional Animal Care and Use Committee (IACUC) under protocol number 2009-0117. All experiments and methods were performed in accordance with relevant guidelines and regulations.

### Cell culture

Adipose stromal cells (ASCs) were isolated from subcutaneous fat of 10-week-old B6.Cg-Lepob/J (*ob*/*ob*) mice and their age-matched C57BL/6J wild-type (WT) controls (Jackson Laboratories) according to previously published protocols^32^. MCF10AT1 cells were obtained from the Barbara Ann Karmanos Cancer Institute. (For the purposes of the companion study reported in Ref. ^31^) MCF10AT1 cells were transfected with a commercially available turbo-green fluorescent protein (GFP) vector (Thermo). Successfully transfected GFP+ cells were sorted on a BD FACS Aria cytometer.

ASCs were cultured in DMEM/F12 media supplemented with 10% fetal bovine serum and 100 U/mL penicillin-streptomycin. MCF10AT1 cells were cultured in enriched DMEM/F12 media supplemented with 5% horse serum, 10 μg/mL insulin, 0.5 μg/mL hydrocortisone, 100 ng/mL cholera toxin, 20 ng/mL EGF, and 100 U/mL penicillin-streptomycin. All cell lines were maintained in incubators at 37°C and 5% CO_2_, with media changes every two days. For co-culture experiments, a 1:1 ratio of the respective media for each cell type was used.

### Spheroid/sample preparation

96-well tissue culture plates were coated with 50 μL/well of 1.5% agarose diluted in DMEM/F12. This coating solidified to form a non-adherent surface (which promotes the coalescence of cells into spheroids). MCF10AT1 cells, ASCs, or co-cultures of the two (in a 1:1 ratio) were seeded into each well of the agarose-coated plate and placed on a rotating shaker (60 rpm) overnight to form multicellular spheroids.

Glass-bottomed petri dishes (Matsunami, 35 mm diameter well) were divided into three distinct wells/sectors via the addition of a handcrafted divider. This divider was made from a 1.5 mm thick sheet of PDMS with three equal-sized compartments cut out, and was covalently bonded to the glass-bottomed petri dishes following plasma treatment.

High concentration rat tail collagen I (Corning) was reconstituted and neutralized with sodium hydroxide (NaOH) and 10× DMEM/F12 to a final concentration of 6 mg/mL. Collagen and spheroids were deposited on the prepared 3-well dishes (1 spheroid/well). In order to promote the formation of thick collagen fibers, the dishes were held at 4°C for 15 min, 20°C for 15 min, and finally 37°C for 15 min. Each gel was then immersed with media and transported to the incubating bio-chamber (described below).

### Imaging system

All images were acquired using a custom-built spectral domain OCM imaging system (depicted in Fig. S1 in Supplementary Note 1) which was developed based upon our previously reported system^30^ with minor modifications, including a novel sample arm design (detailed below and in Supplementary Notes 1 and 2). Light was supplied by a Ti:Sapph laser (Femtolasers, INTEGRAL Element, central wavelength=800 nm, bandwidth=160 nm) and distributed within the microscope by a fiber coupler (Thorlabs, TW805R2A2, 90% to the reference arm, 10% the to sample arm). (The power incident upon biological samples was approximately 4-5 mW.) Polarization in each arm was controlled and matched via manual fiber polarization controllers (Thorlabs, FPC560). Custom-length fiber patch cords were added to reduce the total dispersion mismatch between the two arms.

In the sample arm, light exiting the optical fiber was collimated using a (Thorlabs, AC127-019-B) lens. Beam angle was scanned using a pre-mounted galvanometer mirror pair (Cambridge Technologies, ProSeries I, 10 mm aperture, 10 V analog communication, S4 coating). The separation of the two galvanometer mirrors along the optical axis (as determined by their housing) was *d*=13.69 mm. The two galvanometer mirrors were (approximately) imaged to the back focal plane of the objective lens (Olympus, LCPlan N 20×/0.45 IR, air immersion) via a novel custom-built 1:1 magnification 4F telescope. The telescope consisted of three lenses: two *f*=100 mm lenses (Thorlabs, AC508-100-B) at the front and back, and one *f_x_*=∞, *f_y_*=+700 mm cylindrical lens (Thorlabs, LJ1836L1-B) at the center. The central cylindrical lens was used to compensate for coherence gate curvature (CGC)^30,46^, a distortion artifact which emerges due to the separation (*d*) of the galvanometer mirrors along the optical axis. A theoretical model and results of a validation experiment for this novel OCM system design are provided in Supplementary Notes 1 and 2, respectively. As detailed in Supplementary Note 1, the ‘optimal’ focal length of the cylindrical lens was predicted to be approximately (100 mm)^2^/(13.69 mm)≈+730 mm. The +700 mm lens that was used here was the nearest readily available ‘off-the-shelf’ part. With this design, our system reduced CGC to at most 6% (and may be capable of reducing CGC to as little as 1.5%) of its original severity (i.e., when no cylindrical lens is present).

In the reference arm, light was collimated as in the sample arm and then reflected via a retro-reflector (Thorlabs, PS975M-B) mounted to a single-axis micrometer stage (Newport 9064-X, for fine position adjustment), which was further mounted to a rail (for coarse position adjustment). The amount of returning light was controlled via a manual tunable aperture.

The objective lens was mounted so as to image samples in an inverted configuration (i.e., from underneath). Samples were held above the objective lens in an incubating bio-chamber (Okolab, UNO-PLUS), which was used to maintain physiological temperature, humidity, and pH throughout time-lapse experiments. The bio-chamber was mounted to a 3-axis micrometer stage (Newport, 9064-XYZ-R). Linear stepper motors (Thorlabs, ZST225B motor, KST101 controller, and KCH601 controller hub/power supply) were used to control the position of all 4 micrometer stages in the system (3 axes to control the position of the bio-chamber in the sample arm, and 1 axis to control the distance to the retro-reflector in the reference arm). These motors were controlled via commands issued in MATLAB R2017a.

Raw image data were acquired using a spectrometer (Wasatch Photonics, Cobra 800) with a 2048-pixel line scan camera (Teledyne e2v, Octoplus). Galvanometer mirror scanning and data acquisition were controlled via custom software built in LabView. Data was acquired using a line scan rate of 55 kHz and an exposure time of 10 μs. The system exhibited a sensitivity of ~90 dB and a fall-off of −5 dB/mm. The axial and lateral resolutions of the system were approximately 2.4 μm and 1.5 μm, respectively. In order to perform automated multi-day time-lapse imaging, the data acquisition software, motors, laser, and computer-to-server data transfers were orchestrated via a master control program implemented in MATLAB R2017a.

### Time-lapse imaging protocol

Immediately after spheroid embedding, the sample dish (containing 2-3 spheroids) was placed within the bio-chamber. Manual adjustment of the 4 linear motors was used to establish a set of control points for each spheroid. These control points defined a fixed position for each motor to hold when acquiring images of the corresponding spheroid. The control points for a given spheroid were selected such that: 1) the spheroid was centered within the lateral FOV of the system (sample arm *x/y* control points), 2) the ‘equator’ of the spheroid was roughly aligned to the focal plane (sample arm *z* control point), and 3) the spheroid and coverslip surface (at the base of the dish) appeared within an acceptable range of locations within the axial FOV of the imaging system (reference arm *z* control point). Establishing control points for all of the spheroids in a given dish required approximately 10-15 minutes.

Prior to beginning time-lapse imaging, a ‘calibration image’ was acquired for each spheroid. For a given spheroid, the motors were commanded to move to their corresponding control points. Then, one of the lateral (*x*/*y*) motors was adjusted such that the FOV was centered at a point 1-1.25 mm to the side of the spheroid. This resulted in the spheroid being removed from the FOV, such that only ‘empty’ collagen substrate remained. An image of this region was then acquired for use during OCT image reconstruction. (The use of this calibration image is detailed in Supplementary Note 3, ‘OCT image reconstruction procedure: Focal plane curvature removal’.) This imaging procedure (which was performed manually for all of the spheroids in a given dish) required approximately 5-10 minutes.

Once the above procedures were completed, time-lapse imaging was begun. Images of each spheroid in the dish were acquired at 40-minute time intervals for a total of 48 hours. (In between rounds of imaging, the laser shutter was closed to avoid optical heating of the sample.) For a given spheroid and time-point, nine volumetric images were acquired in rapid succession, similar to the ‘burst protocol’ used in our previous TF-OCM study^30^. The nine images consisted of 1 ‘full FOV’ volume (spanning a lateral FOV of 1.25×1.25 mm^2^ for larger co-culture spheroids, or 1×1 mm^2^ for smaller monoculture spheroids, using 1024×1024 lateral pixels) and 8 ‘reduced FOV’ volumes (spanning a lateral FOV of 937.5×937.5 mm^2^ for larger co-culture spheroids, or 0.75×0.75 mm^2^ for smaller monoculture spheroids, using 768×768 lateral pixels). Each such ‘burst’ of volumetric data generated approximately 24 GB of raw data (resulting in each time-series experiment generating approximately 1.75 TB of raw data per spheroid). Each burst required ~2 minutes to acquire. Imaging of 3 spheroids in a single dish (including sample repositioning between spheroids) required approximately 8-10 minutes per time-point. This left approximately 30 minutes of ‘down time’ between each (40-minute interval) time-point for raw data to be transferred to a remote high-capacity server.

### OCT image reconstruction

Volumetric OCT images were reconstructed from raw image data using a custom procedure similar to that described in our previous study^30^. In brief, this procedure was designed to minimize both spatial distortions within individual OCT volumes and relative spatial distortions *between* OCT images in a single time-series. The procedure consisted of 6 key components: (1) (Depth-selective) OCT volume reconstruction, (2) coherence gate curvature removal, (3) phase registration, (4) focal plane curvature mitigation, (5) bulk demodulation, and (6) computational adaptive optics. (Evidence and discussions regarding the need/reasoning for these procedures may be found in our previous publication^30^.) Since the implementation of these procedures has been updated/modified since our previous study, flow charts and equations describing these updated procedures have been provided in Supplementary Note 3 (‘OCT image reconstruction procedure’). All OCT image reconstruction and subsequent image processing was performed in MATLAB R2018a.

### Drift correction

Despite the use of precision motors for sample positioning, small amounts of bulk drift (along *x, y*, and/or *z*) were present between time-points. The OCT image reconstruction procedures (described in Supplementary Note 3) automatically corrected for drift along the *z*-axis. Drift along the lateral dimensions was measured and corrected separately after OCT image reconstruction.

In order to measure lateral drift, first, an average OCT image was computed via a ‘mean projection’ of the 8 ‘reduced FOV’ images obtained for a given time-point. That is, denoting the *N*=8 reduced FOV images by *S_i_* (*x*, *y*, *z*, *t*) for *i* ∈ {1,2, …, *N*}, the average (‘mean projection’) OCT image 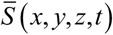 was obtained via:

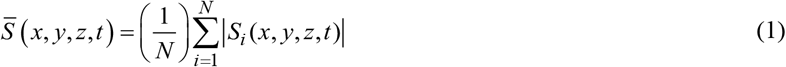

For a given time-point, 4 sub-volumes were extracted from this average OCT image, corresponding to regions along each of the 4 sides of the lateral FOV. The lateral dimensions of these sub-volumes were selected to be as large as possible while still excluding the glass surface and spheroid body (which appeared at the center of the FOV) and thus only contained signals from the surrounding collagen substrate. (For a visual representation, these regions of interest are analogous to the four sides of the region depicted in Fig. S9 in Supplementary Note 3.) The drift of each sub-volume with respect to the corresponding sub-volume of the *previous* time-point was computed to sub-pixel precision via 3D cross-correlation. The total lateral drift of a given time-point with respect to the *previous* time-point was taken to be the median value of the individual drift values reported across all 4 sub-volumes. Total drift with respect to the *first* time-point was then computed via cumulative summation from time *t*=0 up to the given time-point. This lateral drift was removed from all OCT images which were used for subsequent processing.

### Cell and collagen channel synthesis

Reconstructed OCT images were split into two synthetic ‘channels’ corresponding to cells and collagen/background medium, respectively. To achieve this, the 8 reduced FOV images acquired for a given spheroid and time-point were combined via a ‘standard deviation projection’. That is, denoting the *N*=8 reduced FOV images by *S_i_* (*x*, *y*, *z*, *t*) for *i* ∈ {1,2, …, *N*}, the output (‘standard deviation projection’) OCT image *S_σ_* (*x*, *y*, *z*, *t*) was obtained via:

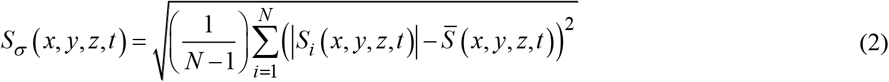

where 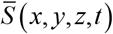 is defined as in Eqn. (1). Similar to our previously reported method^30^, this technique leverages temporal speckle contrast to distinguish rapidly changing structures (primarily cells) from static background structures (here, primarily collagen), resulting in a synthetic ‘cell’ channel. ‘Cell’ channel images were down-sampled by a factor of 768:400 along the lateral dimensions, and by a suitable factor along the axial dimension, via the MATLAB function *imresize3* to yield a final voxel size of (1.88 μm)^3^ and (2.34 μm)^3^ for images of monoculture spheroids and co-culture spheroids, respectively. This down-sampling was used in order to perform smoothing and reduce computational load. These images exhibited a depth-dependent additive ‘background’. This depth-dependent background profile was estimated by computing the median depth-dependent intensity profile from a volumetric region along the periphery of the lateral FOV of the ‘cell’ channel image (analogous to the previously described sub-volumes used for performing lateral drift correction). This background profile was then subtracted from the original image, yielding the completed ‘cell’ channel image.

Preliminary binary images were generated from each (down-sampled) ‘cell’ channel image by applying a 5 ×5×5 voxel median filter followed by thresholding via Otsu’s method. Noise and non-cell structures in these preliminary binary images were mitigated via 4D region growing. Specifically, at the first time-point, only the spheroid (the largest connected structure) was retained from the preliminary binary image. The resulting ‘clean’ binary image then served as the ‘seed’ image for the second time-point. The only structures retained from the preliminary binary image of the second time-point were those which exhibited partial or complete spatial overlap with the structure present in the seed image. This resulted in a clean binary image for the second time-point, which then served as a seed image for the third time-point, and so on. (This algorithm is vulnerable to ‘losing’ cells which break away from the spheroid body and migrate away fast enough as to exhibit no spatial overlap with cells in the image acquired at the previous time-point. However, we did not observe substantial instances of this phenomenon with our 40-minute imaging interval. Decreasing this interval could be used to mitigate this problem in the future.)

In order to generate the ‘collagen’ channel, the volumes in the image sequence 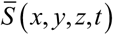 from Eqn. (1) were down-sampled in the same manner as the ‘cell’ channel images described above. The values of any voxels corresponding to ‘cells’ (as determined by the cleaned binary images described above) were set to 0, yielding the complete ‘collagen’ channel image.

### Collagen displacement tracking

Time-varying collagen displacements were computed across the first 24 hours of images using elastic image registration via the built-in MATLAB function *imregdemons* (which uses Thirion’s ‘Demons Algorithm’, an iterative, large-deformation registration method inspired by the optical flow equations^37,38^). Specifically, this function was used to compute the 3D displacement field required to register the ‘collagen’ channel image of the first time-point (*t*=0) to the ‘collagen’ channel image of any given time-point acquired during the first 24 hours of imaging. The *imregdemons* function was called using the following input parameters (as defined on the documentation page for the function^47^): pyramid levels = 3, iterations per pyramid level = 100, and accumulated field smoothing = 2. With these parameters, the resulting displacement measurements exhibited a noise floor of 34 nm for images of monoculture spheroids, and 41 nm for images of co-culture spheroids. Further details and definitions regarding these measurements are provided in Supplementary Note 4.

Displacements were not computed for later time-points beyond the 24 hour mark because the progression of invasive protrusions (and associated collagen degradation and deformation) became substantial, and so the assumptions underlying the Demons Algorithm (and most other standard elastic image registration algorithms) break down^38^. As detailed in the Discussion, further work will be required in the future to obtain deformation tracking results that are sufficiently accurate/reliable for quantitative CTF reconstructions. However, the methods used here have been assumed to be sufficiently accurate to demonstrate the value of our computational 4D-OCM imaging methods for mechanobiology research and future developments in 4D TF-OCM.

### Figure/video generation

All figure/video panels were generated/rendered using MATLAB R2019b. Figures were arranged using Microsoft PowerPoint 2016.

## Acknowledgements

The work described here was supported by funding from the National Institutes of Health through award numbers R01GM132823 and U54CA210184.

## Author contributions

J.A.M. developed and implemented the computational 4D-OCM imaging system, image reconstruction algorithms, and data processing routines. J.A.M. performed all data acquisition and data processing. L.L. performed all cell culture and spheroid/sample preparation. L.L. assisted in data acquisition and analysis of results. C.F. and S.G.A. provided guidance, feedback, and support throughout this study. J.A.M. generated all figures and wrote the manuscript. All authors edited the manuscript.

## Data availability

All relevant data are available from the authors.

## Code availability

All relevant code is available from the authors.

