## Supplementary Information for "Computational 4D-OCM for label-free imaging of collective cell invasion and force-mediated deformations in collagen"

### Contents:

**Supplementary Note 1**: Hardware-based compensation of coherence gate curvature (Model)

**Supplementary Note 2**: Hardware-based compensation of coherence gate curvature (Validation)

**Supplementary Note 3**: OCT image reconstruction procedure

**Supplementary Note 4**: Displacement tracking noise floor

**Supplementary Video Captions**

**Supplementary Note 1: Hardware-based compensation of coherence gate curvature (Model)**

In our previous study[1], we used a fully computational approach to mitigate coherence gate curvature (CGC) artifacts which distort OCT/OCM image data. However, our approach relied on heuristics which only approximated the necessary distortion corrections in order to maintain computational efficiency. In this study, we employed a hybrid CGC mitigation method which physically corrected the majority of CGC via modified optical hardware and employed computational methods to correct for only small residual distortions. This section provides a model for our novel hardware-based mitigation method, while the next section provides results from a validation experiment.

Our microscope (depicted in Fig. S1) is a beam-scanning spectral domain OCM system. Similar to many research- and commercial-grade systems, our microscope uses a set of paired galvanometer mirrors to control scanning of the imaging beam across the lateral FOV. The use of paired galvanometers offers multiple benefits for OCM systems, including simple construction/alignment, compact design, and lower aberrations/losses (due to the reduction or elimination of relay optics ubiquitous to microscopes that use separate/distinct galvanometer mirrors). However, as is shown below, this simple and common 'paired galvanometer' convention in OCM system design can contribute substantial distortions (CGC) to 3D OCT/OCM images when using a wide lateral FOV (e.g., 1×1 mm$^2$ and greater). Thankfully, this fault of the paired galvanometer convention is easily compensated by our novel design, which incorporates a weak cylindrical lens into a 4F telescope relay. This cylindrical lens allows our system to maintain the benefits of a compact and simple paired galvanometer design, while simultaneously mitigating CGC artifacts.

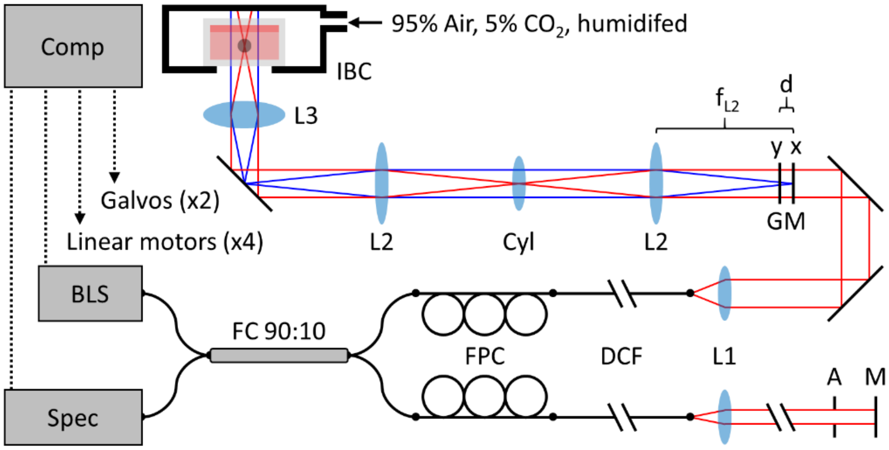


**Figure S1.** Diagram of our novel custom-built OCM imaging system for performing high-resolution, wide-FOV, low-distortion imaging (see the Methods in the main text for a detailed description of components). Comp: computer. BLS: broadband laser source. Spec: spectrometer + line scan camera. FC: fiber coupler (90% to reference arm, 10% to sample arm). FPC: fiber polarization controller. DCF: dispersion compensating fiber (each arm contains different lengths of fiber). A: aperture. M: mirror/retro-reflector. GM: galvanometer mirror (*x*,*y* denote the axis along which each mirror tilts). (Note: Only the *position* of the galvanometer mirrors *along the optical path* is depicted here.) d: galvanometer mirror separation (13.69 mm, imposed by housing). L1: Collimating lens ($f_{L1}$=19 mm). L2: Telescope lens ($f_{L2}$=100 mm). Cyl: Cylindrical lens ($f_x=\infty$, $f_y$=+700 mm, which helps to compensate for coherence gate curvature resulting from the physical separation between the *x* and *y* galvanometers along the optical axis). L3: Objective lens (idealized). IBC: incubating bio-chamber.

Here, we use ray transfer matrix analysis to investigate a simplified model of the sample arm of our OCM imaging system. Within the *xz*-plane, the 'chief' (i.e., central) ray of the collimated beam is parametrized by the position-angle vector $\langle r=0,\theta=0\rangle$. Upon reflection off of the *xz*-plane galvanometer mirror ('$GM_x$'), this ray is imparted with an angle of $\theta_x$ to become $\langle 0,\theta_x\rangle$. Propagating this angled ray through the sample arm up to a distance *z* after the objective lens ('Obj') results in the transformation:

$$\underbrace{\begin{bmatrix}1 & z\\0 & 1\end{bmatrix}}_{\text{Obj}\to z}\underbrace{\begin{bmatrix}1 & 0\\-1/f_o & 1\end{bmatrix}}_{\text{Obj}}\underbrace{\begin{bmatrix}1 & f+f_o\\0 & 1\end{bmatrix}}_{\text{TL2}\to\text{Obj}}\underbrace{\begin{bmatrix}1 & 0\\-1/f & 1\end{bmatrix}}_{\text{TL2}}\underbrace{\begin{bmatrix}1 & f\\0 & 1\end{bmatrix}}_{\text{Cyl}\to\text{TL2}}\underbrace{\begin{bmatrix}1 & 0\\-1/f_x & 1\end{bmatrix}}_{\text{Cyl}}\underbrace{\begin{bmatrix}1 & f\\0 & 1\end{bmatrix}}_{\text{TL1}\to\text{Cyl}}\underbrace{\begin{bmatrix}1 & 0\\-1/f & 1\end{bmatrix}}_{\text{TL1}}\underbrace{\begin{bmatrix}1 & f\\0 & 1\end{bmatrix}}_{\text{GM}_x\to\text{TL1}}\begin{bmatrix}0\\\theta_x\end{bmatrix} \quad \text{(A.1a)}$$

$$\begin{bmatrix}(1/f_o)(z-f_o) & -f_o+\left(f^2/f_o f_x\right)(z-f_o)\\(1/f_o) & \left(f^2/f_o f_x\right)\end{bmatrix}\begin{bmatrix}0\\\theta_x\end{bmatrix} \quad \text{(A.1b)}$$

$$\begin{bmatrix}(1/f_o)(z-f_o) & -f_o\\(1/f_o) & 0\end{bmatrix}\begin{bmatrix}0\\\theta_x\end{bmatrix}, \text{ for } f_x=\infty \quad \text{(A.1c)}$$

$$\begin{bmatrix}-f_o\theta_x\\0\end{bmatrix} \quad \text{(A.1d)}$$

where $f$ denotes the focal length of the telescope lenses (‘TL1’ and ‘TL2’), $f_x$ denotes the focal length of the cylindrical lens (‘Cyl’) within the *xz*-plane, and $f_o$ denotes the focal length of the (idealized) objective lens. Since $GM_x$ is imaged to the back focal plane of the objective lens, the beam is scanned in an ideal telecentric fashion (i.e., the beam remains parallel to the optical axis while its lateral *x*-position is scanned proportionally to $\theta_x$ ). Of course, deviations from this idealized model (e.g., thick lenses, aberrations in the telescope and/or objective, etc.) will cause non-ideal beam-scanning behavior at large scanning angles/lateral beam positions.

We further analyze the optical path length (OPL) traversed by the chief ray through the system. In the small-angle limit, propagation through air by a distance $L$ (along the optical axis) at an angle $\theta$ results in an OPL accumulation of approximately $L\left(1+\theta^2/2\right)$. Likewise, transmission through a thin lens of focal length $f$ at a lateral position $r$ with respect to the optical axis accumulates an OPL (up to an additive constant) of approximately $-r^2/2f$ . Following the path of the chief ray within the *xz*-plane, we derive a total accumulated OPL (up to an additive constant) of:

$$\begin{aligned}\text{OPL}_x(z,\theta_x) = {} & \underbrace{f\left(1+\theta_x{}^2/2\right)}_{\text{GM}_x\to\text{TL1}}-\underbrace{(f\theta_x)^2/2f}_{\text{TL1}}+\underbrace{f}_{\text{TL1}\to\text{Cyl}}-\underbrace{0}_{\text{Cyl},\,f_x=\infty}+\underbrace{f}_{\text{Cyl}\to\text{TL2}}-\underbrace{(f\theta_x)^2/2f}_{\text{TL2}}+\ldots\\ & \underbrace{(f+f_o)\left(1+(-\theta_x)^2/2\right)}_{\text{TL2}\to\text{Obj}}-\underbrace{(-f_o\theta_x)^2/2f_o}_{\text{Obj}}+\underbrace{z}_{\text{Obj}\to z}+\text{constant}\end{aligned} \quad \text{(A.2a)}$$

$$\text{OPL}_x(z,\theta_x)=4f+f_o+z+\text{constant} \quad \text{(A.2b)}$$

This corresponds to an ideal scenario, since OPL increases linearly with *z* and is not a function of scanning angle.

Within the *yz*-plane, we perform a similar analysis. The chief ray is imparted with an angle $\theta_y$ by the *yz*-plane galvanometer mirror (‘$GM_y$’) to become $\left\langle 0,\theta_y\right\rangle$. Propagation of this angled ray through the sample arm up to a distance *z* after the objective lens results in the transformation:

$$\underbrace{\begin{bmatrix}1 & z\\0 & 1\end{bmatrix}}_{\text{Obj}\to z}\underbrace{\begin{bmatrix}1 & 0\\-1/f_o & 1\end{bmatrix}}_{\text{Obj}}\underbrace{\begin{bmatrix}1 & f+f_o\\0 & 1\end{bmatrix}}_{\text{TL2}\to\text{Obj}}\underbrace{\begin{bmatrix}1 & 0\\-1/f & 1\end{bmatrix}}_{\text{TL2}}\underbrace{\begin{bmatrix}1 & f\\0 & 1\end{bmatrix}}_{\text{Cyl}\to\text{TL2}}\underbrace{\begin{bmatrix}1 & 0\\-1/f_y & 1\end{bmatrix}}_{\text{Cyl}}\underbrace{\begin{bmatrix}1 & f\\0 & 1\end{bmatrix}}_{\text{TL1}\to\text{Cyl}}\underbrace{\begin{bmatrix}1 & 0\\-1/f & 1\end{bmatrix}}_{\text{TL1}}\underbrace{\begin{bmatrix}1 & f-d\\0 & 1\end{bmatrix}}_{\text{GM}_y\to\text{TL1}}\begin{bmatrix}0\\\theta_y\end{bmatrix} \quad \text{(A.3a)}$$

$$\begin{bmatrix}(1/f_o)(z-f_o) & -f_o+\left(\left(f^2-f_y d\right)/f_o f_y\right)(z-f_o)\\(1/f_o) & \left(\left(f^2-f_y d\right)/f_o f_y\right)\end{bmatrix}\begin{bmatrix}0\\\theta_y\end{bmatrix} \quad \text{(A.3b)}$$

$$\begin{bmatrix}-f_o\theta_y+(z-f_o)\alpha\theta_y\\\alpha\theta_y\end{bmatrix}, \text{ for } \alpha=\left(\left(f^2-f_y d\right)/f_o f_y\right) \quad \text{(A.3c)}$$

where $d$ denotes the separation between $GM_x$ and $GM_y$ along the optical axis. Here, since $GM_y$ is *not* imaged to the back focal plane of the objective lens, the beam does *not* scan parallel to the optical axis (i.e., the output beam propagates at an angle which varies as a function of $\theta_y$). Moreover, the lateral $y$-position of the beam varies as a function of both $\theta_y$ and depth $z$. These behaviors result in one of the hallmarks of CGC: a depth-dependent lateral magnification (i.e., non-telecentricity).

Likewise, following the path of the chief ray within the $yz$-plane, we derive a total accumulated OPL (up to an additive constant) of:

$$\begin{aligned}\mathrm{OPL}_y\left(z,\theta_y\right) = &\underbrace{d}_{\mathrm{GM}_x\to\mathrm{GM}_y} + \underbrace{(f-d)\left(1+\theta_y^{\,2}/2\right)}_{\mathrm{GM}_y\to\mathrm{TL1}} - \underbrace{\left((f-d)\theta_y\right)^2/2f}_{\mathrm{TL1}} + \underbrace{f\left(1+\left(d\theta_y/f\right)^2/2\right)}_{\mathrm{TL1}\to\mathrm{Cyl}} - \underbrace{\left(f\theta_y\right)^2/2f_y}_{\mathrm{Cyl},\,f_y} + \ldots \\ &\underbrace{f\left(1+\left(\left(d/f - f/f_y\right)\theta_y\right)^2/2\right)}_{\mathrm{Cyl}\to\mathrm{TL2}} - \underbrace{\left(\left(f+d-f^2/f_y\right)\theta_y\right)^2/2f}_{\mathrm{TL2}} + \underbrace{\left(f+f_o\right)\left(1+\left(-\theta_y\right)^2/2\right)}_{\mathrm{TL2}\to\mathrm{Obj}} + \ldots \\ &-\underbrace{\left(\left(-f_o+d-f^2/f_y\right)\theta_y\right)^2/2f_o}_{\mathrm{Obj}} + \underbrace{z\left(1+\left(\left(-d/f_o+f^2/f_o f_y\right)\theta_y\right)^2/2\right)}_{\mathrm{Obj}\to z} + \text{constant}\end{aligned} \tag{A.4a}$$

$$\mathrm{OPL}_y\left(z,\theta_y\right) = \left[4f + f_o + z\right] + \left[-\alpha f_o + \left(z - f_o\right)\alpha^2\right]\theta_y^{\,2}/2 + \text{constant} \tag{A.4b}$$

where $\alpha$ is defined as in Eqn. (A.3c). Here, we observe that OPL varies quadratically as a function of $\theta_y$. This results in the namesake 'curvature' of CGC. That is, flat surfaces will appear curved in the OCT image due to the fact that OCT microscopes use OPL measurements as a proxy for 'depth'.

Given this analysis, we hypothesized that CGC could be mitigated (within the confines of this idealized model) if $f_y = f^2/d$. This scenario would result in $\alpha = 0$, and would thus eliminate the quadratic terms in Eqn. (A.4b). In our system, the optimal focal length for our cylindrical lens was predicted to be $f_y$=(100 mm)$^2$/(13.69 mm)≈+730 mm. However, since we used a non-optimal focal length of only +700 mm, our system was not expected to completely eliminate CGC. However, compared to the case of applying no correction (i.e., when $f_y$=+∞), our system was *predicted* to reduce CGC (i.e., $\alpha$) to approximately 5% of its original severity (i.e., when no compensation method is employed).

**Supplementary Note 2: Hardware-based compensation of coherence gate curvature (Validation)**

In order to assess the performance of our CGC-compensating system, we acquired images of a flat glass surface near the focal plane of our system, with and without the CGC-compensating cylindrical lens and spanning a 1×1 $mm^2$ lateral FOV. The resulting images (and apparent curvature of the glass surface) are depicted in Fig. S2. We then fit each surface (hereafter denoted by indices $i$=1 and $i$=2, respectively) with a quadratic polynomial:

$$z_i(x,y) = a_i x^2 + b_i xy + c_i y^2 + d_i x + e_i y + f_i \quad \text{(A.5)}$$

The terms $a_i$, $b_i$, and $c_i$ record the apparent curvature of the glass surface and thus measure the severity of CGC. The terms $d_i$ and $e_i$ record the apparent tilt of the glass surface, which results from a combination of true sample tilt and/or non-ideal beam alignment within the sample arm optics (the latter of which can be compensated with an adjustable periscope placed prior to the galvanometers). The term $f_i$ encodes the axial offset of the glass surface in the image.

The curvature terms for the imaged glass surfaces were found to have values (in units of $\mu m^{-1}$) of $a_1 = 4.585\times10^{-5}$, $b_1 = 0.015\times10^{-5}$, and $c_1 = -4.280\times10^{-5}$ for the uncorrected system, and $a_2 = -1.454\times10^{-6}$, $b_2 = 0.856\times10^{-6}$, and $c_2 = -2.415\times10^{-6}$ for the CGC-compensated system. In order to assess the total amount of CGC, we performed a rotation of the lateral coordinate system to make the cross-term $b_i$=0. Specifically, defining:

$$\begin{bmatrix} x \\ y \end{bmatrix} = \begin{bmatrix} \cos\theta_i & \sin\theta_i \\ -\sin\theta_i & \cos\theta_i \end{bmatrix} \begin{bmatrix} x' \\ y' \end{bmatrix} \quad \text{(A.6)}$$

we obtained the transformed curvature terms ($A_i$, $B_i$, and $C_i$) via:

$$\begin{bmatrix} A_i \\ B_i \\ C_i \end{bmatrix} = \begin{bmatrix} \cos^2\theta_i & -\cos\theta_i\sin\theta_i & \sin^2\theta_i \\ \sin 2\theta_i & \cos 2\theta_i & -\sin 2\theta_i \\ \sin^2\theta_i & \cos\theta_i\sin\theta_i & \cos^2\theta_i \end{bmatrix} \begin{bmatrix} a_i \\ b_i \\ c_i \end{bmatrix} \quad \text{(A.7)}$$

where $\theta_i = 0.5\arctan\left(b_i/(c_i - a_i)\right)$, yielding $A_1 = 4.585\times10^{-5}$, $B_1 = 0$, and $C_1 = -4.280\times10^{-5}$ for the uncorrected system, and $A_2 = -1.291\times10^{-6}$, $B_2 = 0$, and $C_2 = -2.578\times10^{-6}$ for the CGC-compensated system. The opposite signs of $A_1$ and $C_1$ correspond to positive CGC along one axis and negative CGC along the orthogonal axis, respectively. This corresponds to the scenario where the back focal plane of the objective lens is imaged by the 4F telescope to a plane that lies *between* the two galvanometer mirrors. In contrast, the matching negative signs of $A_2$ and $C_2$ indicate that the alignment of our CGC-compensated system was not optimal (i.e., the objective lens was placed too close to the telescope relay by a small margin). Despite this non-optimal alignment, our system managed to reduce CGC to *at most* $\max\left(|A_2|,|C_2|\right)/\min\left(|A_1|,|C_1|\right) \approx 6\%$ of its original severity, which is close to our theoretically predicted value of 5%. However, under optimal alignment conditions, we would expect to achieve the condition $A_i = -C_i$ for both systems (i.e., the back focal plane of the objective lens is imaged to a plane that lies at the mid-point between the galvanometer mirrors). Thus, defining the 'total' CGC of the system as $|C_i - A_i| = |2A_i| = |2C_i|$ under optimal alignment conditions, we estimate that our system is capable of reducing CGC to approximately $|C_2 - A_2|/|C_1 - A_1| \approx 1.5\%$ of its original severity. This exceeds our predicted improvement and could be attributable to spherical aberration of the telescope lenses causing their effective focal lengths to decrease at large scanning angles (and thus making them better-matched to the +700 mm focal length of the installed cylindrical lens).

Given these results, we conclude that our novel OCM imaging system provides both a simple and effective method to mitigate CGC artifacts which emerge from the use of paired galvanometer mirrors. As a result, our system substantially reduces a potential source of distortions which would otherwise corrupt deformation data that is essential to traction force microscopy. Since it remains difficult to realize an ideal system, computational methods should still be employed to mitigate small residual distortions. Such methods are detailed in the next section of this document.

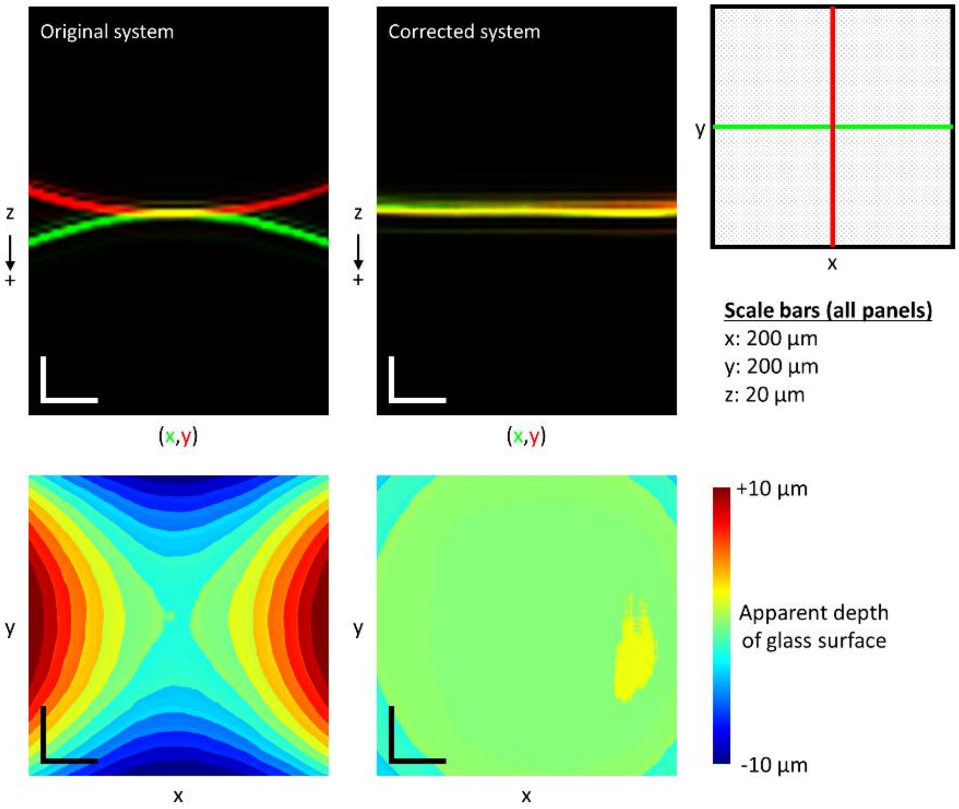


**Figure S2.** Visualization of coherence gate curvature (CGC) before (left) and after (right) hardware-based compensation. The top panels depict the OCM image of a (*physically* flat) glass-air interface in the *xz*- (green) and *yz*- (red) planes intersecting the origin of the lateral FOV. Any *observed* curvature of the surface in these panels is a consequence of CGC distortions. The bottom panels depict the observed apparent depth (i.e., the optical path length) of the surface as a function of lateral position. In the original system, CGC causes image distortions which result in the (physically flat) glass surface appearing as a hyperbolic paraboloid surface. In the corrected system, the glass surface appears nearly flat (with a residual weak paraboloid shape due to non-optimal alignment). See text for details.

**Supplementary Note 3: OCT image reconstruction procedure**

Here, we provide detailed flow charts and equations for our OCT image reconstruction procedure. Note that more detailed explanations/justifications underlying many of the operations below may be found in Supp. Ref. [1].

**Depth-selective OCT volume reconstruction**

OCT images were reconstructed from spectral data by applying the standard operations of background subtraction, spectrum resampling, dispersion compensation, and the Fourier transform. In order to enable the reconstruction of small/specific depth ranges (as opposed to the full depth range of the available axial FOV), a matrix multiplication-based method analogous to previously reported algorithms[2-4] was employed. Specifically, for a (post-background subtraction) A-scan of spectral data $v(k_b, x, y)$ for $b \in \{1, 2, \ldots, 2048\}$, the OCT image signal $s(z_a, x, y)$ at a given integer depth index $a$ was computed via:

$$s(z_a, x, y) = \sum_{b=1}^{2048} v(k_b, x, y) \exp\left(-j\left(2k_b z_a + \alpha_2\left(k_b - \bar{k}\right)^2 + \alpha_3\left(k_b - \bar{k}\right)^3\right)\right) = \sum_{b=1}^{2048} \Theta_{ab} v(k_b, x, y) \qquad \text{(B.1)}$$

where $a$ is constrained to the set of integers $a \in \{1, 2, \ldots, 2048\}$, $j = \sqrt{-1}$, $k_b$ denotes the wavenumber of the $b^{\text{th}}$ spectrometer pixel (determined by manufacturer-generated data sheets), $\bar{k} = (k_1 + k_{2048})/2$, $z_a = \Delta z(a - 1025)$ for $\Delta z = (2047/2048)(\pi / |k_{2048} - k_1|)$, and $(\alpha_2, \alpha_3)$ denote (manually calibrated) dispersion compensation parameters. Note that $\Delta z$ corresponds to the height of each voxel in terms of optical path length (OPL). The physical height of a given voxel is $\Delta z / n$, where $n$ is the local refractive index of the imaged medium.

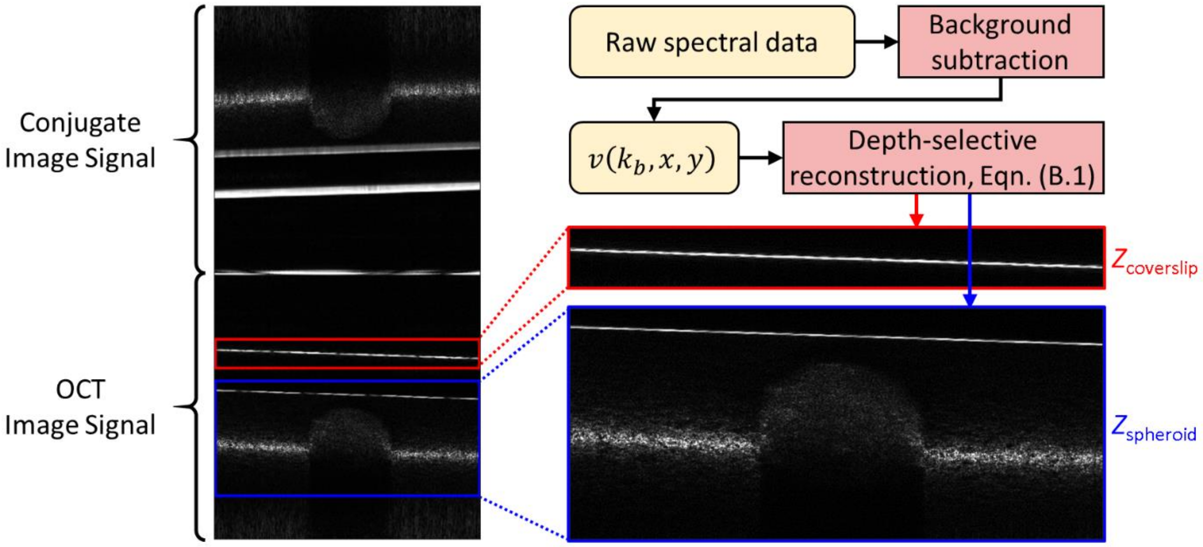


**Figure S3.** Depth-selective OCT image reconstruction and regions of interest. See text for details.

There were two regions of interest in the imaged volumes. The first is a narrow range of depths $z_a \in Z_{\text{coverslip}}$ which contain the coverslip surface (i.e., the base of the sample dish) across all time-points. The second is a larger range of depths $z_a \in Z_{\text{spheroid}}$ which contain the spheroid and surrounding regions across all time-points. Examples of these regions are depicted in Fig. S3.

**Coherence gate curvature removal and phase registration**

As discussed in the previous sections of this supplementary document, coherence gate curvature (CGC) is a consequence of non-idealities in imaging system design/implementation and/or sample positioning which distort OCT images. A flat surface may appear to be tilted and/or curved in OCT images acquired by a system corrupted by CGC. Unlike our previously reported methods[1], which used a single computational procedure for mitigating CGC, here we employed a three-pronged hybrid approach to obtain more robust results. The majority of CGC was removed via the cylindrical lens incorporated into our novel OCM imaging system (discussed previously, and depicted in Fig. S1). Residual CGC was removed computationally in two steps, hereafter called 'coarse' and 'fine' CGC removal. The latter step simultaneously performs phase registration (another key operation for OCT image reconstruction). CGC calibration and removal were performed independently for each image and time-point (except where otherwise noted).

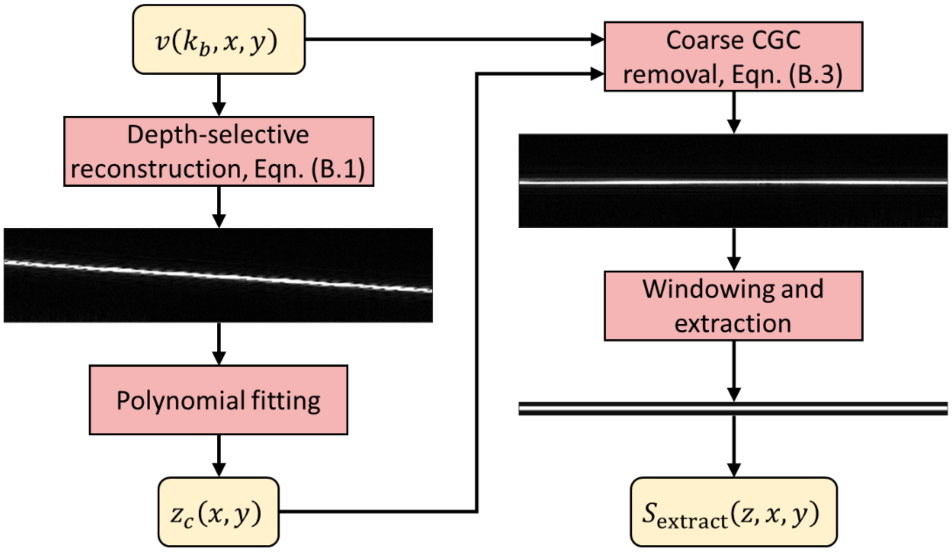


**Figure S4.** Calibration procedure for both 'coarse' (left column) and 'fine' (right column) coherence gate curvature removal (and phase registration). See text for details.

The first step is a calibration stage (depicted in Fig. S4), which performs calibrations for both coarse and fine CGC removal (and phase registration). An OCT image of the $Z_{\text{coverslip}}$ region is reconstructed according to Eqn. (B.1). The laterally varying axial position of the coverslip surface (in terms of OPL) is then approximated by the function:

$$z_c(x, y) = c_{xx}x^2 + c_{xy}xy + c_{yy}y^2 + c_x x + c_y y + c_0 \quad \text{(B.2)}$$

This completes the calibration stage for coarse CGC removal. Next, the $Z_{\text{coverslip}}$ region is reconstructed *again* while applying coarse CGC removal via:

$$\hat{S}(z_a, x, y) = \sum_{b=1}^{2048} \Theta_{ab} v(k_b, x, y) \exp\left(-j2k_b\left(z_c(x, y) - z_0\right)\right) \quad \text{(B.3)}$$

where $z_0$ is defined as the value of $c_0$ obtained for the *first* time-point in the time-lapse data set. In the resulting 3D image, the coverslip will appear *nearly* flat and level, and will be centered at the OPL position $z = z_0$. Note that by using the same value of $z_0$ for all time-points, all reconstructed images in a given time-series will be automatically registered/aligned along the *z*-axis. The coarsely-corrected image is next cropped along the *z*-axis such that only ~11-21 voxels remain along the *z*-axis centered around the position $z = z_0$. This cropped volume will be referred to as $S_{\text{extract}}(z, x, y)$. This completes the calibration stage for fine CGC removal and phase registration.

The cropped volume $S_{\text{extract}}(z, x, y)$ is used to perform both fine CGC removal and phase registration. Note that although this volume was derived from the $Z_{\text{coverslip}}$ region of the sample, it may be applied to the reconstruction of any sub-region from the full OCT data set (e.g., the $Z_{spheroid}$ region). In order to perform fine CGC removal and phase registration to a volume $P(z,x,y)$, $S_{\text{extract}}(z,x,y)$ is first zero-padded along the $z$-axis until it is of the same size as $P(z,x,y)$. Then, the following operation is performed:

$$P_r(z,x,y) = FT^{-1}_{z\to q_z}\left[FT_{z\to q_z}\left[P(z,x,y)\right]\exp\left(-j\arg\left(FT_{z\to q_z}\left[S_{\text{extract}}(z,x,y)\right]\right)\right)\right] \quad \text{(B.4)}$$

where $FT_{z\to q_z}$ and $FT^{-1}_{z\to q_z}$ denote the forward and inverse Fourier transform along $z$, respectively, and $P_r(z,x,y)$ denotes the output volume (which lacks CGC and is phase-registered). This procedure is depicted in Fig. S5.

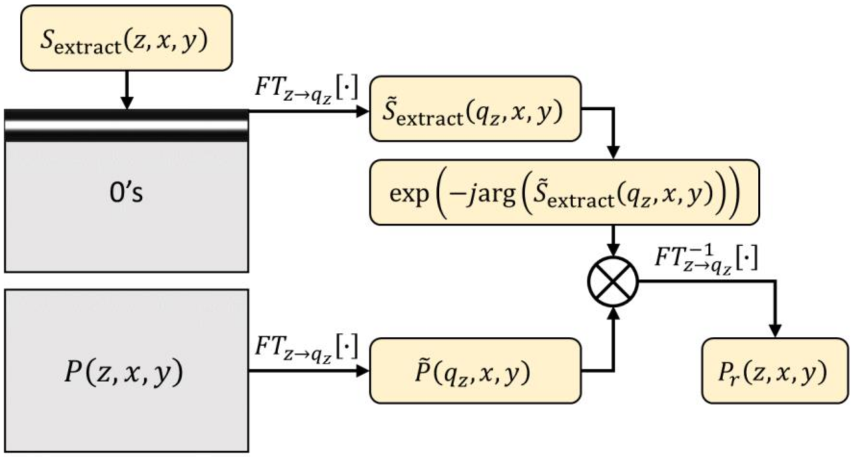


**Figure S5.** Fine coherence gate curvature removal and phase registration. See text for details.

**Focal plane curvature removal**

Focal plane curvature (FPC) is another consequence of non-optimal system design and/or sample positioning. This results in the focal 'plane' appearing tilted and/or curved, even after CGC removal has been performed (i.e., the curvature of the focal plane does not necessarily match that of the 'coherence gate'). This results in OCT images which exhibit a laterally-varying point spread function (PSF), which conventional refocusing algorithms (such as the computational adaptive optics (CAO) procedure used in this study) do not accommodate, due to the assumption of a laterally-invariant PSF. Alternative formulations of CAO which account for lateral variation do exist[5,6]. However, such procedures can be computationally expensive and complicated to calibrate and perform. In our previous study[1], we demonstrated an alternative procedure for FPC mitigation which enables the use of computationally efficient CAO algorithms that leverage lateral invariance assumptions.

Unlike our previous study[1] (which measured/calibrated FPC from images of the sample directly), here we used a separate calibration data set (which is described in the Methods of the main text). This calibration data set was used to obtain a measurement of FPC, which was subsequently applied to all images in the corresponding time-series. The reason for this change in procedure is a consequence of the new experimental settings encountered in this study. In our previous study, we imaged isolated cells which occupied a miniscule fraction of the total volumetric FOV. Therefore, FPC could be readily measured from images of the sample. However, the spheroids imaged in this study occupied a large fraction of the volumetric FOV, and substantially obstructed the focal plane across much of the FOV. This prevented automated measurements of FPC using only images of the spheroid(s). The calibration data set used in this study contained no spheroid (but was instead merely acquired *near* the spheroid). This provided a clear and unobstructed view of the geometry of the focal 'plane' across the entire FOV. The FPC measured from this data set was assumed to be a suitable proxy for FPC calibration of the spheroid images in a given time-series.

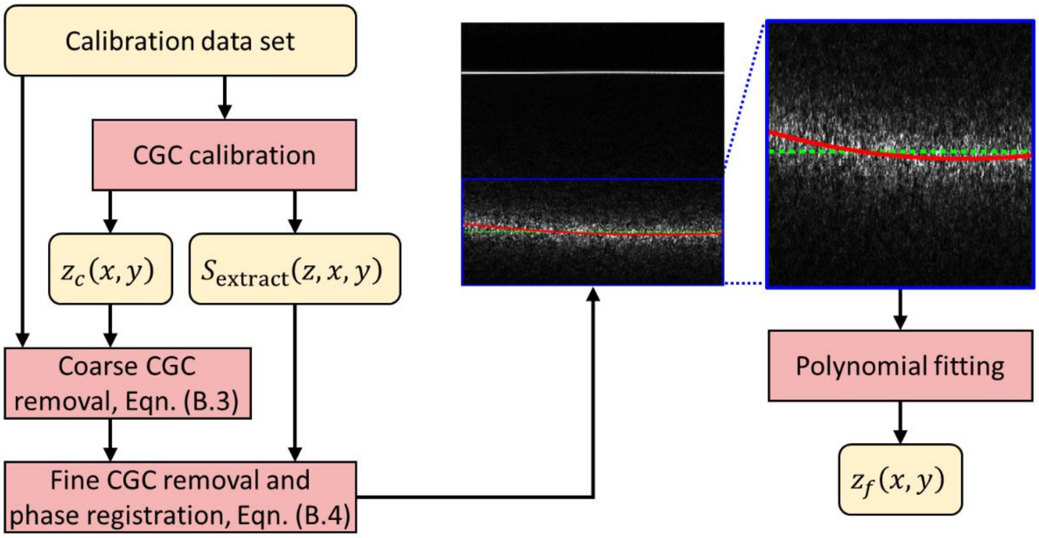


**Figure S6.** Calibration procedure for focal plane curvature removal. Note that the focal plane appears curved even though the coverslip surface appears flat and level. See text for details.

FPC calibration was performed as depicted in Fig. S6. First, CGC calibration is performed on the calibration data set. Then, the 'focal plane region' of the sample (which is analogous to the $Z_{\text{spheroid}}$ region, but with no spheroid present, given the nature of the calibration data set) is reconstructed while applying coarse CGC removal, fine CGC removal, and phase registration. The laterally varying position of the focal 'plane' (in terms of OPL) is approximated via:

$$z_f(x, y) = f_{xx}x^2 + f_{xy}xy + f_{yy}y^2 + f_x x + f_y y + f_0 \tag{B.5}$$

This completes the calibration stage for FPC removal. In order to apply FPC removal (as depicted in Fig. S7), the $Z_{\text{spheroid}}$ region of the spheroid images are first reconstructed via:

$$\hat{S}(z_a, x, y) = \sum_{b=1}^{2048} \Theta_{ab} v(k_b, x, y) \exp\left(-j2k_b\left(z_c(x, y) - z_0\right)\right) \exp\left(-j2k_b\left(z_f(x, y) - \bar{z}_f\right)\right) \tag{B.6}$$

where $\bar{z}_f$ is defined as the mean value of $z_f(x, y)$ across the lateral FOV. Note that this formula is nearly identical to that for coarse CGC removal (Eqn. (B.3)), with an extra phase term added for performing FPC removal as well. Following this reconstruction, fine CGC removal and phase registration are performed via Eqn. (B.4) to obtain a phase registered image in which the focal plane appears flat and level (although the coverslip surface appears curved, due to the mismatch between CGC and FPC). This volume, which we will call '$S_r(z,x,y)$', is compatible with digital refocusing procedures which assume a laterally invariant PSF[1,7].

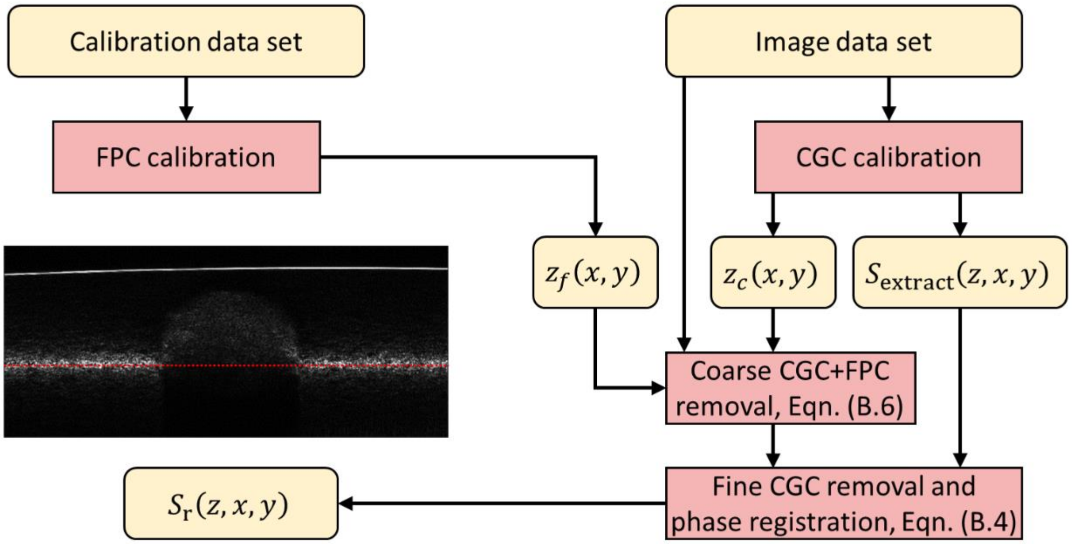


**Figure S7.** Procedure for joint application of coarse CGC removal, fine CGC removal, phase registration, and FPC removal. Note that this results in an image where the focal plane is flat and level, but the coverslip surface is not. See text for details.

**Bulk demodulation**

As discussed in our previous study[1], optical misalignments and/or sample tilt can result in the OCT image signal undergoing a bulk phase modulation across the lateral dimensions. Failing to account for this modulation can result in depth-dependent shearing artifacts which emerge after applying CAO. Calibration of the bulk modulation in the system was performed as depicted in Fig. S8. First the 'full FOV image' (defined in the Methods section of the main text) from the first time-point in a given time-series is reconstructed using the FPC removal procedure detailed previously. Next, any depths containing a glass surface are windowed out from the volume or set to zero (since the signal from such reflective surfaces can overwhelm/corrupt the calibration procedure). The remaining volume then undergoes the following operation:

$$M(q_x, q_y) = \int \left| FT_{3D} \left[ S_r(z,x,y) \right] \right| dq_z \tag{B.7}$$

where $FD_{3D}$ denotes the 3D forward Fourier transform operation. The resulting real-valued function $M(q_x,q_y)$, which is depicted in the upper-right panel of Fig. S8, approximates the lateral spatial frequency content of the image signal. Given the use of Gaussian beams in our system, $M(q_x,q_y)$ has an approximately Gaussian profile and is off-centered with respect to the origin ($q_x$=0,$q_y$=0) due to the bulk phase modulation of the image signal (which we wish to measure and remove). Performing peak-finding along the $q_x$- and $q_y$-axes results in a measurement of the bulk modulation coefficients: ($q_{x,0}$,$q_{y,0}$).

Once this calibration was performed, all volumes in the time-lapse data set were demodulated according to:

$$S_d(z,x,y) = S_r(z,x,y)\exp\left(-j\left(q_{x,0}x + q_{y,0}y\right)\right) \tag{B.8}$$

In doing so, it was assumed that the bulk modulation was constant across the entire time-lapse data set.

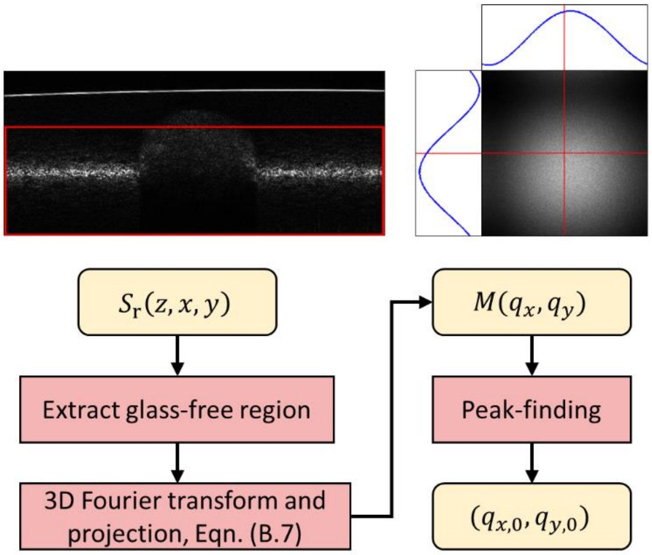


**Figure S8.** Calibration procedure for bulk demodulation. See text for details. The red boxed region in the upper-left panel shows the region that is used to perform the calibration (note that it excludes the glass surface which appears near the top of the image). $M(q_x,q_y)$ is shown in the upper right panel. The intersection of the two red lines denotes the origin of the lateral spatial frequency domain.

**Computational adaptive optics**

Computational adaptive optics[7] (CAO) is used to compensate for depth-dependent degradation of the lateral resolution of the OCT system. Here, we assumed that this degradation was due entirely to defocus (and not higher-order optical aberrations). In order to perform CAO, first the depth of the focal plane must be calibrated. For a given time-point, the 'full FOV image' was reconstructed using the procedures described above (CGC removal, FPC removal, phase registration, and bulk demodulation). Next the average axial intensity profile was computed from the 3D region of interest depicted in Fig. S9. (Note that this region excludes both the spheroid body as well as any glass surfaces which appear in the image.) Curve-fitting was performed in order to find the peak of this axial intensity profile and thus obtain the depth $z_{focus}$ of the focal plane (in terms of physical distance). This position was used for both the full FOV and reduced FOV images of the spheroid (defined in the Methods section of the main text) for the given time-point.

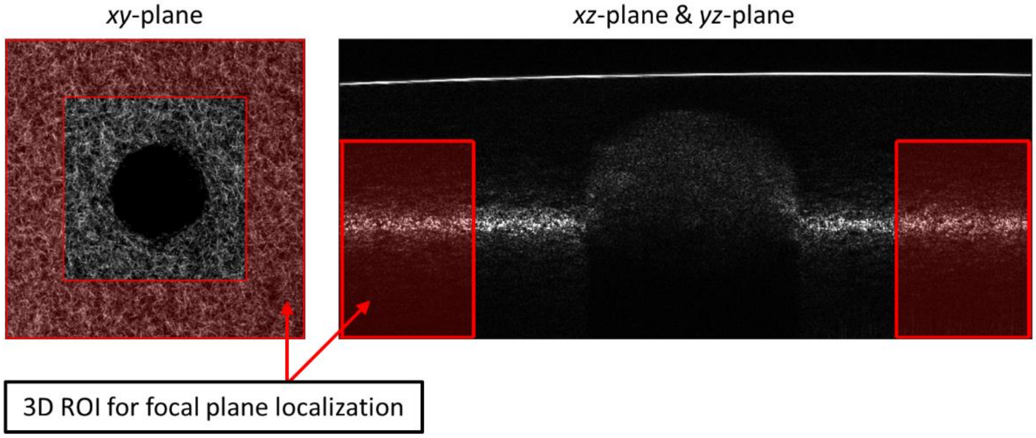


**Figure S9.** 3D region of interest for axial localization of the focal plane. The image shown in the right-hand panel intersects the origin of the lateral FOV. See text for details.

Next, defocus-compensation was performed via:

$$S_f(z,x,y) = FT^{-1}_{(x,y)\to(q_x,q_y)}\left[FT_{(x,y)\to(q_x,q_y)}\left[S_d(z,x,y)\right]\exp\left(-j\left(z-z_{\text{focus}}\right)\sqrt{\left(2n\bar{k}\right)^2-q_x^{\,2}-q_y^{\,2}}\right)\right] \quad \text{(B.9)}$$

where $FT_{(x,y)\to(q_x,q_y)}$ and $FT^{-1}_{(x,y)\to(q_x,q_y)}$ denote the forward and inverse 2D Fourier transform across the lateral dimensions, respectively, $n$ denotes the refractive index of the sample (here, $n$=1.34 was used), and $\bar{k}$ is defined as before (and corresponds to the central wavenumber measured by the spectrometer of imaging system). Following this operation, the magnitude of the (complex-valued) image signal was normalized with respect to $z$. (Note that the normalized image remains complex-valued!) The results of this procedure are depicted in Fig. S10.

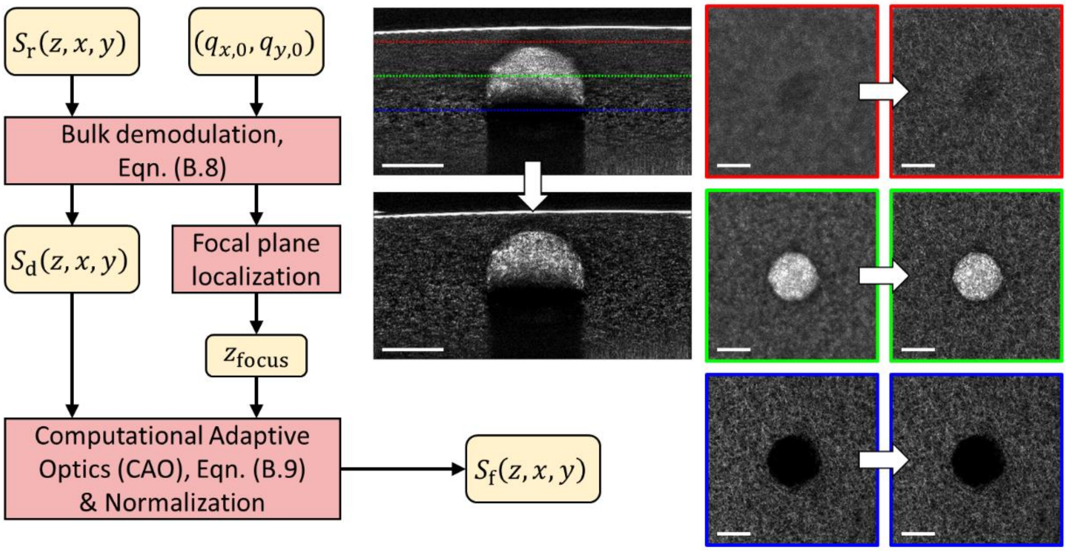


**Figure S10.** Procedure for defocus compensation with computational adaptive optics. See text for details. Panels at the right depict *en face* planes before and after CAO at the depths indicated by the three colored lines spanning the top-middle panel. Red/top: 225 µm above the focal plane. Green/middle: 112.5 µm above the focal plane. Blue/bottom: Focal plane. Scale bars = 200 µm.

**Focal plane curvature restoration**

In this study, we applied the same 'ideal coordinate system' heuristic established in our previous study[1]. That is, the ideal/final coordinate system for our reconstructed OCT images was assumed/defined as the one in which the coverslip surface appears both flat and level. However, CAO could only be performed while the *focal plane* was flat and level. Therefore, the final step in OCT image reconstruction is to *restore* the focal plane curvature to its original state, which returns the coverslip surface to its 'ideal' flat and level state.

For a reconstructed OCT volume with $N \leq 2048$ pixels along the axial dimension (recall that our spectrometer camera had 2048 pixels), a frequency-axis vector $\mathbf{q}_z \in \mathbb{R}^{N\times 1}$ is defined such that:

$$q_{z,i} = \left(\frac{2\pi}{|\Delta z|}\right)\left(\frac{i-1}{N}\right), \text{ for } i \in \{1,2,...,N\} \quad \text{(B.10)}$$

where $\Delta z$ is defined (in terms of OPL) as before. Then, FPC restoration is performed via:

$$FT^{-1}_{z\to q_z}\left[FT_{z\to q_z}\left[S_f(z,x,y)\right]\exp\left(-jq_z\left(z_f(x,y)-\bar{z}_f\right)\right)\right] \quad \text{(B.11)}$$

The final reconstructed image that results from this procedure is depicted in Fig. S11. Note that this volumetric image signal is a complex-valued function. All subsequent image processing (detailed in the Methods section of the main text) were performed on the *magnitude* of this function.

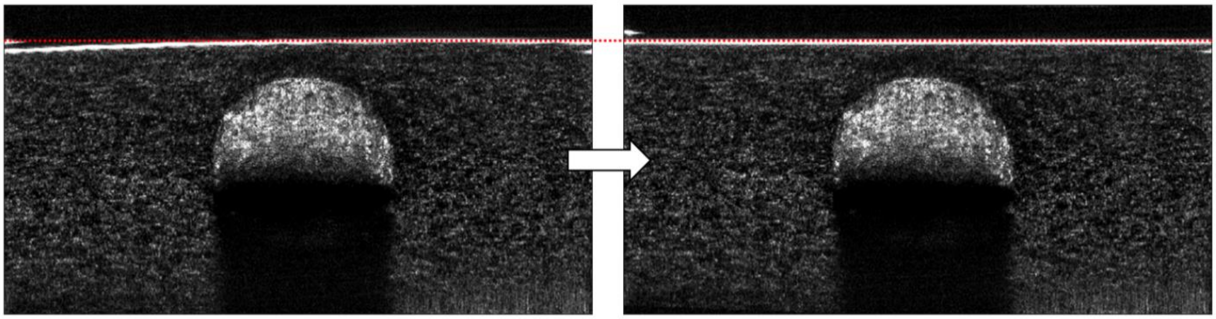

**Figure S11.** Focal plane curvature restoration. The CAO-processed and intensity-normalized image (left) is transformed to a final output image (right) wherein the coverslip surface appears both flat and level. See text for details.

**Supplementary Note 4: Displacement tracking noise floor**

In order to estimate the noise floor of our displacement tracking procedure, we performed a 'self-mapping' test. Our 'burst' imaging protocol (described in the Methods) yielded several redundant images (the 8 'reduced FOV' volumes) of the collagen substrate at each time-point. Due to the short duration over which these images were acquired (less than 2 minutes for a given 'burst'), the observed collagen structure is expected to undergo only negligible deformations. Therefore, if displacement tracking is performed between images obtained from a single time-point, the output displacement field estimate will be dominated by noise artifacts emerging from a combination of imaging system noise and limitations of the tracking algorithm.

To this end, two modified 'mean projections' were obtained (analogous to Eqn. (1) in the Methods) at time *t*=0:

$$\bar{S}_1(x,y,z,t=0)=\left(\frac{1}{4}\right)\sum_{i=1}^{4}\left|S_{2i-1}\left(x,y,z,t=0\right)\right| \quad \text{(C.1)}$$

$$\bar{S}_2(x,y,z,t=0)=\left(\frac{1}{4}\right)\sum_{i=1}^{4}\left|S_{2i}\left(x,y,z,t=0\right)\right| \quad \text{(C.2)}$$

That is, the two 'mean projections' were obtained from non-overlapping subsets of the 8 'reduced FOV' volumes $\{S_1,...,S_8\}$ acquired for the given time-point. (It should be noted that the imaging noise present in both $\bar{S}_1$ and $\bar{S}_2$ is expected to be higher than that of $\bar{S}$ from Eqn. (1) due to each signal being generated from only half the total images used to generate $\bar{S}$. Therefore, the displacement tracking noise floor estimates provided here may be larger than the true displacement tracking noise floor due to larger imaging noise contributions.) The volumes $\bar{S}_1$ and $\bar{S}_2$ were then converted into 'collagen' channel images via the procedure defined in the Methods (i.e., the volumes were subjected to down-sampling, and voxels corresponding to 'cells' were set to 0). Displacement tracking was then performed via the *imregdemons* function (using the same parameters defined in the Methods) to estimate the displacement field:

$$\mathbf{u}(x,y,z)=\left\langle u_x(x,y,z),u_y(x,y,z),u_z(x,y,z)\right\rangle \quad \text{(C.3)}$$

which maps $\bar{S}_1$ to $\bar{S}_2$. The displacement tracking noise floor for each dimension (*x*, *y*, and *z*) was estimated by computing the standard deviation of the associated displacement field components ($u_x$, $u_y$, and $u_z$, respectively) throughout the collagen substrate within a depth range spanning the 'surface' of the spheroid (i.e., the edge closest to the coverslip bottom of the petri dish) to the 'equator' (middle) of the spheroid. These noise floors, $(\sigma_x,\sigma_y,\sigma_z)$ were computed for 6 ASC monoculture spheroids, and 6 ASC + MCF10AT1 co-culture spheroids. The average noise floors computed across these samples were (38 nm, 35 nm, 30 nm) for the monoculture spheroids and (43 nm, 42 nm, 39 nm) for the co-culture spheroids. The average noise floor $\sigma$ (reported in the Methods) was defined as $\sigma=(\sigma_x+\sigma_y+\sigma_z)/3$, yielding σ = 34 nm and σ = 41 nm for the monoculture and co-culture cases, respectively. Normalizing $\sigma$ by the (down-sampled, isotropic) voxel side-length of the images used for displacement tracking (1.88 μm for monoculture spheroids and 2.34 μm for co-culture spheroids, as described in the Methods), yields a displacement tracking noise floor of approximately 0.02 voxels for both cases.

**Supplementary Video Captions**

**Supplementary Video 1**: Animation of the spheroid depicted in Figure 1. Panels (top-left, top-middle, top-right, bottom-left, and bottom-middle) correspond to rows 1-5 from Figure 1, respectively.

**Supplementary Video 2**: Animation of the spheroid depicted in Figure 2. Panels (top-left, top-middle, top-right, bottom-left, and bottom-middle) correspond to rows 1-5 from Figure 2, respectively.

**Supplementary Video 3**: Animation of the panels depicted in Figure 3. The left-hand side depicts panels with only scattering contrast. The right-hand side depicts the same panels including superimposed temporal speckle contrast-based labeling in green (i.e., as shown in Figure 3).

**Supplementary Video 4**: Animation of Figure 4. Time-stamps shown are in HH:MM format.
